# Evolution of a large and diverse phospholipase gene cluster that defines the plant pathogenic genus *Ceratocystis*

**DOI:** 10.64898/2026.07.28.741319

**Authors:** C. G. Mayers, K.S. Kim, M.A. Ferreira, T.C. Harrington

**Affiliations:** Plant Pathology and Plant-Microbe Biology Section, School of Integrative Plant Science, Cornell University, Ithaca, New York 14850; Department of Plant Pathology, Entomology and Microbiology, Iowa State University, Ames, Iowa 50011; Department of Plant Pathology, Universidade Federal de Lavras, Lavras, Brazil 37203-202

## Abstract

Many *Ceratocystis* species cause cankers and unique vascular wilt diseases, often on a broad and unpredictable range of plant hosts. Characteristic necrosis of xylem parenchyma cells and dark staining of surrounding tissue is typically evident, especially in woody hosts. The molecular basis for this unique pathogenicity and host range remains unclear, but bacterial-type phosphatidylinositol phospholipase C (bPI-PLC) genes were recently identified in unusually high copy number in multiple *Ceratocystis* species, and the PLCs may play a role in host membrane disruption. We produced a high-quality long-read genome assembly of the rapid ʻŌhi’a death pathogen, *Ceratocystis lukuohia*, and identified 81 partial or complete PLC-like genes, each with a unique DNA sequence, encoding signal peptides and a PLC-X domain. The putative translations mostly ranged from 300 to 500 amino acids that differed markedly from the fungal and prokaryotic bPI-PLCs at sites conferring phosphatidylinositol specificity, suggesting a novel family of secreted PLCs (Cer-PLCs). Remarkably, 73 of the full or partial Cer-PLC genes reside in a single 543 kb gene cluster in *C. lukuohia*. Comparison to an available long-read genome assembly of *C. fimbriata* revealed a similar Cer-PLC cluster of 61 genes, with a gene order and arrangement broadly similar to that of the *C. lukuohia* cluster, except for a large inversion at the beginning of the cluster. Differences suggest that the cluster is dynamic, with many apparent indels involving multiple Cer-PLCs. We compared 40 newly-assembled genomes of *Ceratocystis* strains and eight publicly available genomes and found that the Cer-PLCs comprise a gene family present in all *Ceratocystis* species but differing greatly in number (26 to 92), with 64 to 92 in species of the highly aggressive Latin American Clade. The two closest relatives of *Ceratocystis* have Cer-PLCs but not in the gene cluster: *Chalaropsis* spp. have only one Cer-PLC, and *Berkeleyomyces basicola* has 25 related Cer-PLCs scattered across multiple contigs. No Cer-PLC was detected in the more-distant members of the Ceratocystidaceae. The unique cluster in *Ceratocystis* apparently arose through insertion of Cer-PLCs within an ancestral gene cluster with a CeGAL transcription factor, followed by repeated duplications and rapid diversification of Cer-PLCs, perhaps driven by unequal crossover events. This extraordinary expansion, diversification, and maintenance of Cer-PLCs may have played a major role in the evolution of aggressiveness and host range in *Ceratocystis*.

**Impact Statement:** New strains of *Ceratocystis* species with expanding host ranges are emerging as important plant pathogens around the world. However, little is known about the basis for the wide variation in host range and aggressiveness of *Ceratocystis* species. An earlier study had identified a gene family coding for phosphatidylinositol-specific phospholipase C (PI-PLC) in some *Ceratocystis* species. Our sequence analyses suggest that the coded enzyme is not likely phosphatidylinositol-specific but may have retained capability of degrading plant membranes and may be a major determinant of aggressiveness and host range. The most aggressive species in the genus has up to 92 copies of this unique class of PLCs, defined here as Cer-PLCs, making the expansion of this gene family among the largest known in fungi. Most of the Cer-PLC genes occur in a gene cluster of more than 500 kb, which appears to be under the control of a CeGAL-type transcription factor. Coordinated regulatory control may enable the hyper-production of these membrane-degrading enzymes during pathogenesis. The Cer-PLC gene family occurs in close relatives of *Ceratocystis*, but the Cer-PLC gene cluster is unique and universal in *Ceratocystis*. The gene cluster is the largest known for a single gene family, and it is highly dynamic and likely undergoes frequent recombination. Multiple introductions of strains to a new environment could generate very aggressive recombinants that attack previously unrecognized hosts, as appears to be happening with the multiple introductions of the South American species *C. manginecans* to Asia.

## INTRODUCTION

*Ceratocystis* species are wound colonizers and plant vascular pathogens, with the most aggressive pathogens causing wilt and mortality of trees (Harrington et al. 2024). Some of the species are specialized to a host genus or family, but many of the species have broad host ranges across many unrelated families of plants. *Ceratocystis fimbriata* on *Ipomoea*, *C. platani* on *Platanus*, and *C. cacaofunesta* on *Theobroma* and *Herrania* are examples of host-specialized species (Baker et al. 2003, Engelbrecht and Harrington 2005, Harrington et al. 2024). However, the *Ipomoea* strain of *C. fimbriata* was recently shown to infect other, distantly-related hosts (Avila et al. 2026), and many *Ceratocystis* species (eg. *C. manginecans*) attack a wide range of unrelated plant families (Harrington et al. 2024). Even closely-related strains of a *Ceratocystis* species can exhibit widely different host ranges (Baker et al. 2003; Harrington et al. 2011, 2024; Valdetaro et al. 2019). A few species, including the well-known *C. fimbriata sensu stricto,* cause dry, black rots of corms or storage roots of herbaceous plants (Thorpe et al. 2005, Li et al. 2016), but the majority of *Ceratocystis* species are pathogens of woody hosts (Harrington 2013). They may colonize xylem and phloem tissues around fresh wounds and cause cankers, but in the most economically important diseases, they move systemically in the xylem and cause lethal vascular wilt diseases (Harrington 2013, Harrington et al. 2024, Hughes et al. 2020). In contrast to “true” vascular wilt pathogens, which move systemically in host xylem before parenchyma cells are killed (Dimond 1970), *Ceratocystis* species typically induce a black stain in colonized xylem, especially along ray parenchyma tissues, and may kill cambium and inner bark (secondary phloem) tissue, causing a canker-stain disease (Harrington 2013, DeBeer et al. 2014, Hughes et al. 2020). The crowns of affected trees may show wilt symptoms as the necrosis of the vascular tissues of branches and stems progress, though large trees may take years to be fully colonized and die (Harrington et al. 2013).

The underlying mechanisms for the unique symptomatology and varying host ranges of species and strains of *Ceratocystis* have been proposed but not conclusively demonstrated. A hydrophobin discovered in *C. platani*, ceratoplatanin, was hypothesized to be a general *Ceratocystis* phytotoxin or effector protein (Pazzagli et al. 1999; Comparini et al. 2009), but homologs of the gene have since been found in a wide range of both pathogenic and non-pathogenic fungi (Gao et al. 2020). Genomic studies have identified various genes coding for potential effector proteins and other pathogenicity factors, though their role in pathogenesis has not been demonstrated (Van der Nest et al. 2015; Zhang et al. 2020; Fourie et al. 2019, 2020; Ramos-Lizardo et al. 2023, Maguvu et al. 2023). Parada-Rojas et al. (2024) predicted 188 effector proteins in a *C. fimbriata* genome assembly (Stahr et al. 2024), and 31 effectors were expressed early in the infection cycle. Fourie et al. (2019) found two quantitative trait loci (QTLs) in *C. fimbriata* associated with aggressiveness to *Ipomoea* and one QTL in *C. manginecans* associated with aggressiveness to *Acacia,* and numerous genes of interest were identified in these regions. A proteomic study of two strains of *C. manginecans* infecting *Acacia* and *Eucalyptus* (Syazwan et al. 2025) identified proteins associated with DNA repair and cellular respiration in the early stages of infection, whereas host-specificity was associated with reprogramming of metabolism and modulation of cellular transport. Cong et al. (2023) identified an APSES transcription factor (Swi6) downstream from the cell wall integrity regulatory (CWI) pathway in *C. fimbriata* attacking *Ipomoea*; deletion of Swi6 greatly reduced virulence. Deletion of (CWI)-mitogen-activated protein kinase (MAPK) genes resulted in defects in hyphopodia formation, which appeared to be needed for leaf infection, and greatly reduced virulence (Lu et al. 2025). Araújo et al. (2024) studied the volatile compounds of *C. cacaofunesta* and found among them a potentially phytotoxic 3-methylbutan-1-ol.

Molano et al. (2018) reported unexpectedly high copy numbers of a bacterial-like phosphatidylinositol-specific phospholipase C (phosphatidylinositol diacylglycerol-lyase; plcA; EC 4.6.1.13; hereafter “bPI-PLC”) in the genomes of *C. cacaofunesta* and four other *Ceratocystis* species. Many of the bPI-PLCs genes coded for N-terminal secretory signal peptides, and four were identified in the *C. cacaofunesta* secretome, suggesting that bPI-PLCs may act as extracellular pathogenicity factors. Later studies confirmed high bPI-PLC copy number in *C. fimbriata* and *C. destructans* (Fourie et al. 2020, Maguvu et al, 2023).

The bPI-PLCs found in bacteria differ from eukaryotic PI-PLCs (EC 3.1.4.11; hereafter “ePI-PLC”) typically found in fungi and other eukaryotes (Heinz et al. 1998; Feng et al. 2003; Macrae et al. 2025). Both hydrolyze phosphatidylinositol and share a catalytic “X domain” with analogous active and catalytic sites (Heinz et al. 1998; Gellatly et al. 2012). The ePI-PLCs are mostly known for their role in signal transduction and have additional domains, a more complex tertiary structure, and a requirement for Ca^2+^ cofactors (Williams and Katan 1996; Katan and Williams 1997; Katan and Cockcroft 2020; Fang et al. 2023; Macrae et al. 2025). The simpler bPI-PLCs have only the single X domain, are often secreted extracellularly, and can cleave glycosylphosphatidylinositol-anchored proteins from eukaryotic cell surfaces, potentially interfering with host immunity (Heinz et al. 1998; Griffith et al. 1999; Roberts et al. 2018).

The PI-PLC genes discovered by Molano et al. (2018) in *Ceratocystis cacaofunesta* have more similarity to bPI-PLCs than to ePI-PLCs. They hypothesized that the *Ceratocystis* bPI-PLCs were phosphatidylinositol-specific and functioned analogously to bacterial bPI-PLCs, either by interacting with membrane-bound molecules or by disrupting membrane integrity and inducing cell death (Molano et al. 2018). Such disruption could explain the unique parenchyma necrosis and vascular staining that is characteristic of *Ceratocystis* diseases (Harrington 2013, DeBeer et al. 2014, Hughes et al. 2020).

*Ceratocystis lukuohia* causes a lethal vascular wilt disease on the native Hawaiian tree *Metrosideros polymorpha*, ʻŌhi’a lehua (Barnes et al. 2018; Cannon et al. 2022; Hughes et al. 2020). The pathogen was apparently introduced to Hawaii. Its closest relatives are *C. xanthosomatis*, *C. platani*, and *C. costaricensis*, which are believed native to the Caribbean region and eastern USA (Harrington et al. 2024). Typical of *Ceratocystis* species in the Latin American Clade, *C. lukuohia* infects wounds and systemically colonizes stems and branches, eventually causing death (Hughes et al. 2020). Infected ʻŌhi’a trees show extensive red reaction zones in the woody xylem and black staining that typically follows along the ray parenchyma. Necrosis and staining of the secondary phloem is also common where in contact with xylem staining. Another *Ceratocystis* species, *C. huliohia*, is not systemic and causes only limited cankering on ʻŌhi’a trees (Juzwik et al. 2024). Though this species appears to be of much less importance than *C. lukuohia*, it has been considered to be a second cause of rapid ʻŌhi’a death (Barnes et al. 2018; Cannon et al. 2022).

The first aim of this study was to develop a near-complete genome assembly of *Ceratocystis lukuohia* using long reads (PacBio) in order to identify bPI-PLC genes and test the hypothesis that the genes occur in a gene cluster (Molano et al. 2018) or on a dispensable chromosome (Mehrabi et al. 2017) as in, for example, *Alternaria alternata* HST clusters (Harimoto et al. 2007; Tsuge et al. 2013). We used publicly available genomes and generated new assemblies of *Ceratocystis* species and their relatives in the family Ceratocystidaceae to determine the phylogenetic relationships within the family and infer the evolutionary history of the bPI-PLCs genes and their genomic organization. We also compared the putative amino acid sequences of the *Ceratocystis* bPI-PLC to those of other eukaryotic and bacterial PLCs to see if they were unique to *Ceratocystis* or the family.

## MATERIALS AND METHODS

### Short read genome sequencing and assembly

Genomic DNA from 49 isolates of *Ceratocystis*, *Chalaropsis*, and *Berkeleyomyces* (Table 1) was extracted from pure cultures grown on malt yeast extract agar for 5–7 d with the Wizard Genomic DNA Purification Kit (Promega). Illumina library prep (NEBNext Ultra) and MiSeq sequencing (2 x 300 bp) was performed by the Iowa State University (ISU) DNA Facility. Reads were cleaned of adapters and quality-trimmed in Geneious Pro v. 11.1 (Biomatters) using BBDuk 1.0 (quality ≥ 13, both ends), then assembled *de novo* using SPAdes 3.11.1 (Nurk et al. 2013, Prjibelski et al. 2020) with default paired-end parameters. Short (< 500 bp) and low-coverage (<33.33% of median contig coverage) contigs were removed, as were low-GC contigs identified as mitochondrial via local BLAST against the full mitogenome of six *Ceratocystis* species (Mayers et al. 2021) or identified as plasmids by NCBI BLASTn.

**Table 1.**
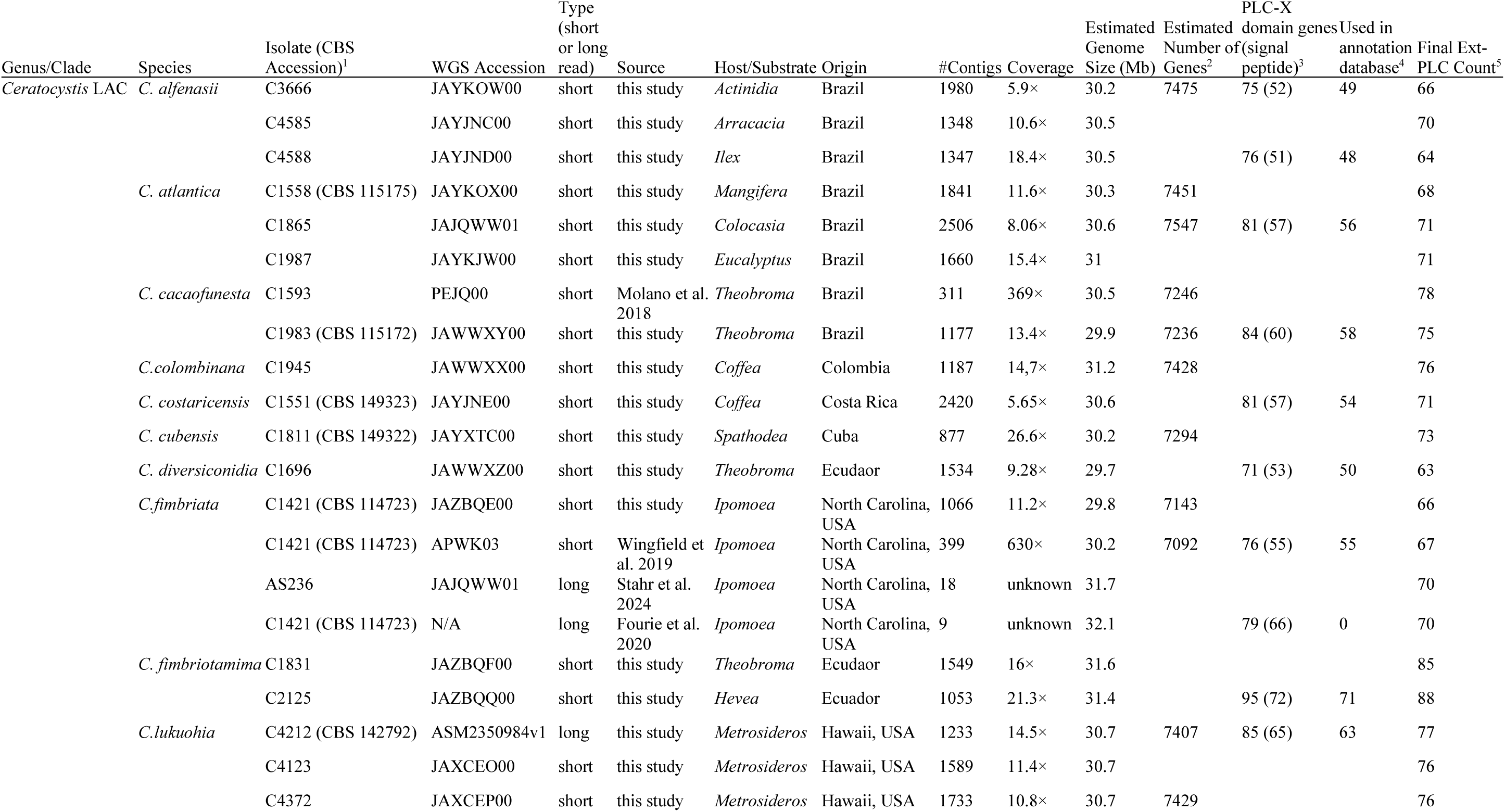
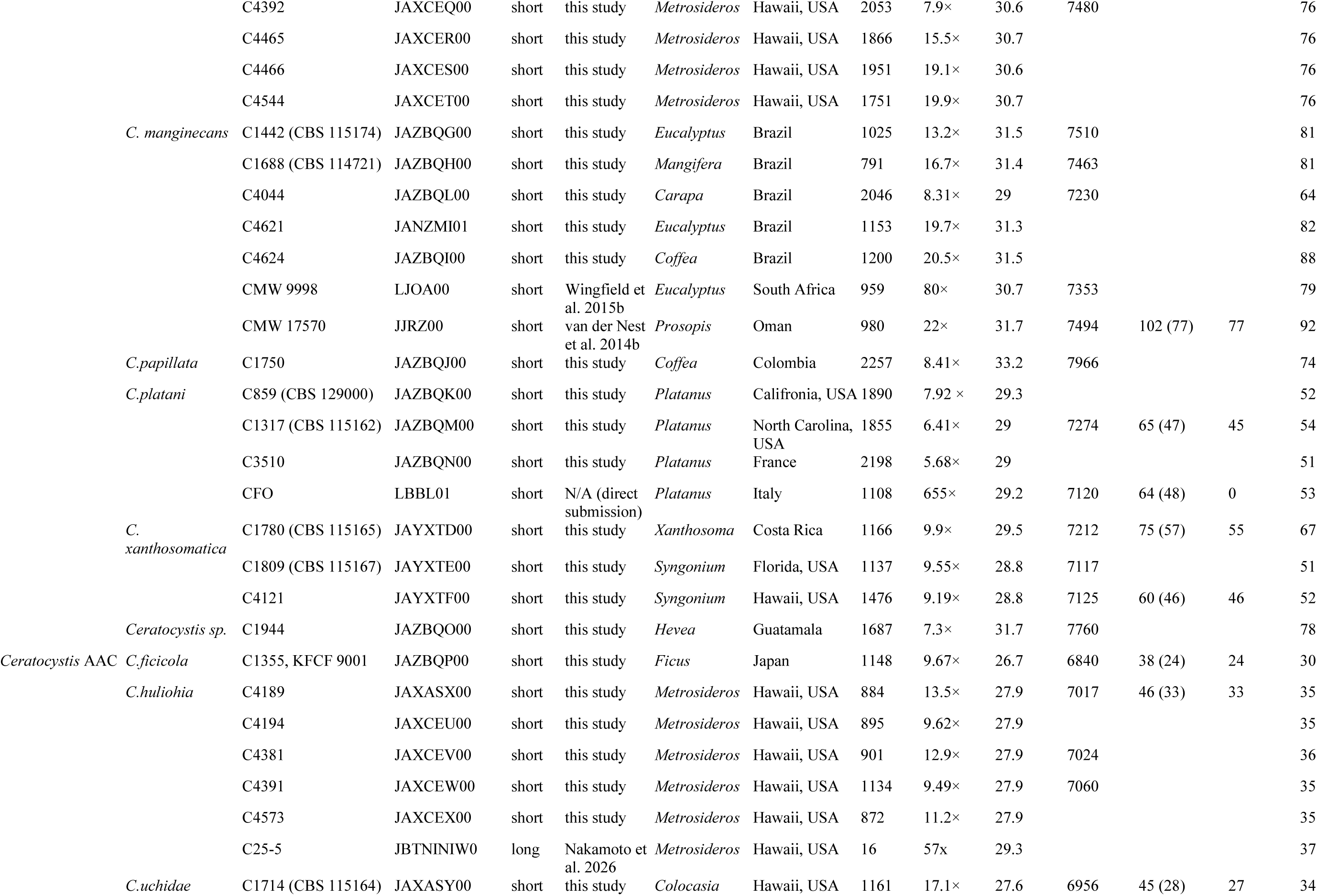
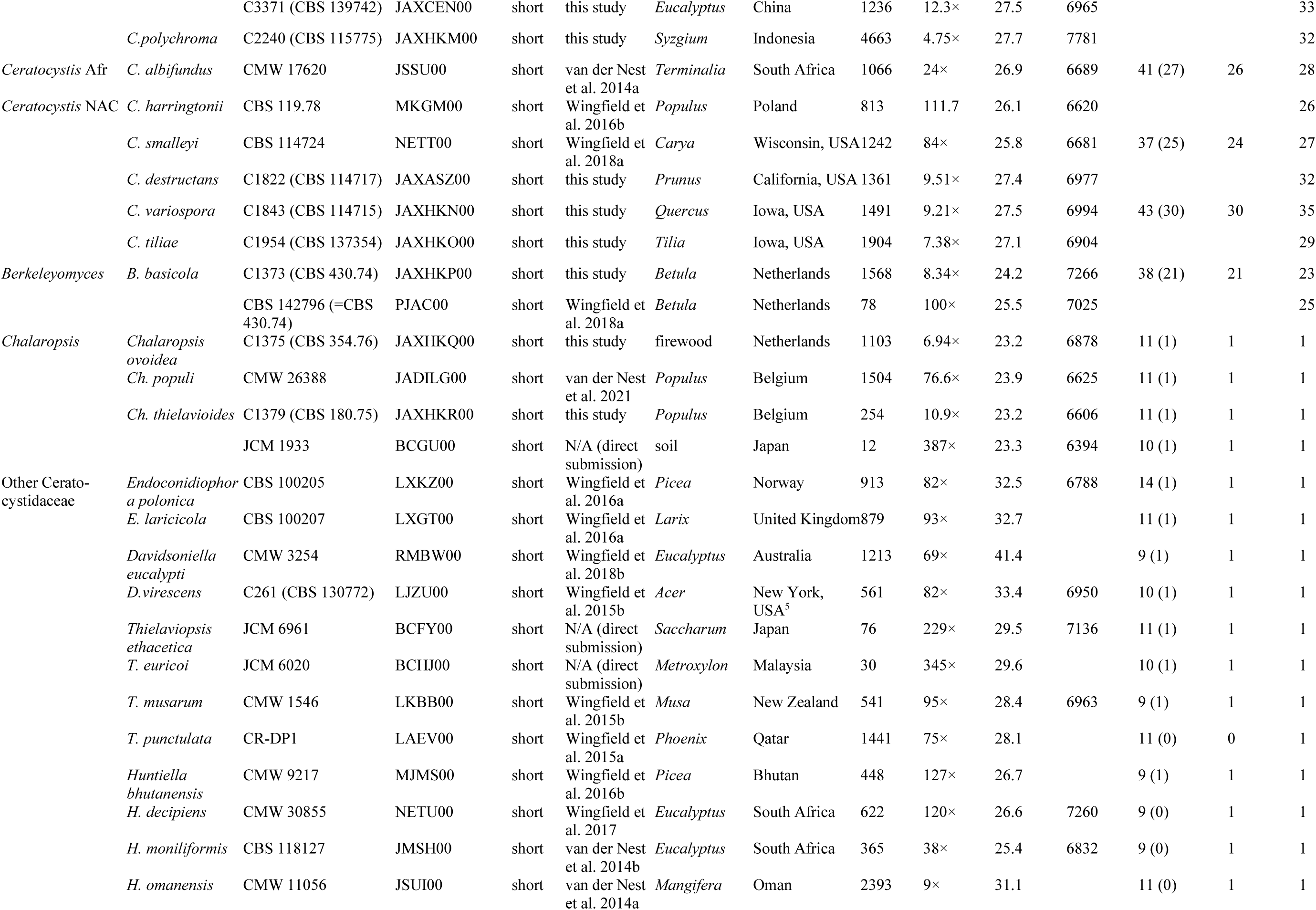
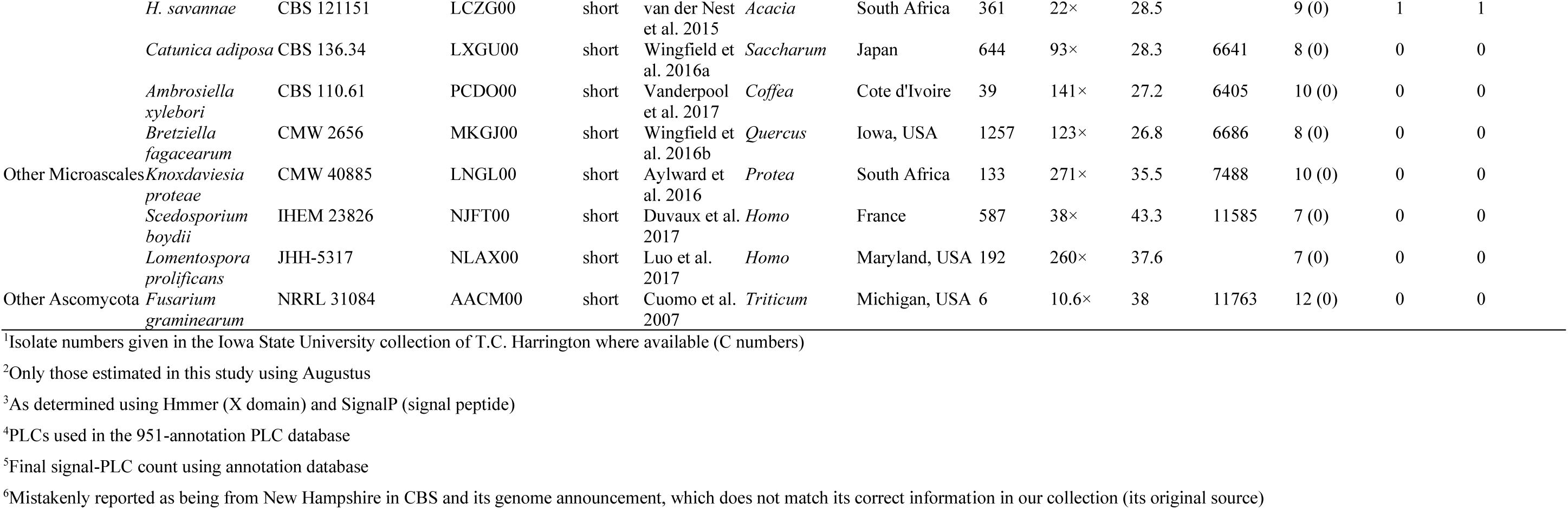
Genome sources and statistics and estimates of PI-PLCs for species of the Latin American Clade (LAC), Asian-Australasian Clade (AAC), African Clade (Afr), and North American Clade (NAC) of *Ceratocystis*, other Ceratocystidaceae and Ascomycota relatives.

### Long read genome sequencing and assembly

Single Molecule, Real-Time (SMRT) sequencing was performed for *C. lukuohia* ex-holotype isolate C4212 (CBS 142792) grown in liquid malt-yeast extract for 5–7 d then vacuum-filtered (Whatman Grade 1, Cytiva). Extracted genomic DNA (E.Z.N.A Fungal DNA Mini Kit, Omega Bio-tek) was quantified (Qubit 2.0, Invitrogen), and 20 µg used to prepare 20 kb SMRTbell libraries sequenced on four PacBio RS II SMRT cell at the ISU DNA Facility. After read correction with default settings (PacBio SMRT Portal v. 2.0.0, Pacific Biosciences, Menlo Park, California), 865,971 subreads were filtered to 361,519 (mean read length 5.88 kb, N50 7.34 kb, 66.38× coverage), which were *de novo* assembled (the “HGAP assembly”) with default parameters in HGAP4 (Chin et al. 2013) and polished (variant-calling) with Quiver (SMRT Link 7.0.0, Pacific Biosciences). A second, independent long-read assembly (the “Canu assembly”) was generated using Canu v. 1.7.1 (Berlin et al. 2015), with default parameters and error rate = 0.045, then polished with Pilon v. 2 (Walker et al. 2014). A third “SPAdes hybrid assembly” was constructed by combining the long-read PacBio RS II sequences with Illumina paired-end reads using hybridSPAdes (Antipov et al. 2016) in SPAdes with default parameters. Assembly completeness was evaluated using BUSCO v. 3.0.1 with the Sordariomycetes odb9 dataset (Simão et al. 2015; Waterhouse et al. 2018).

### Whole genome annotation

Genes were predicted with AUGUSTUS v. 3.3.3 via WebAUGUSTUS (Hoff and Stanke 2013) using the *Fusarium graminearum* training set (Stanke et al., 2005). Selected gene predictions were refined by manual BLASTp, InterProScan (Jones et al. 2014), and MOTIFsearch (Pfam, NCBI-CDD, and PROSITE) searches. Secretion and subcellular localization of translated proteins were assessed with SignalP 6.0 (Teufel et al. 2022) and DeepLoc 2.0 (Thumuluri et al. 2022).

Candidate bacterial-like PI-PLC genes were identified by screening the amino acid translations of the predicted ORFs (Geneious Pro v. R11.1) for the bPI-PLC-like X domain (Pfam PF00388.19) using HMMER 3.2.1 (Mistry et al. 2013) and the Pfam 32.0 protein database (El-Gebali et al. 2019), then checked against the UniProtKB/Swiss-Prot protein database using BLASTp. Those with signal peptides (SignalP 6.0) were classified (DeepLoc 2.0) as putatively secreted bPI-PLCs, or “Ext-bPI-PLCs”; those with cytoplasmic localization as “Cyt-bPI-PLCs”; and those with vacuolar/lysosomal localization as “Vac-bPI-PLCs”.

Functional annotation of predicted proteins used BLASTp against NCBI nr (E≤ 1e-3, 40% identity) via EBlast2GO (Conesa et al. 2005) as implemented in OmicsBox 1.2.4 (BioBam Bioinformatics S.L.), KEGG KO assignment via BlastKOALA v. 2.2 (Kanehisa et al. 2016), ortholog grouping via OrthoMCL v. 2 as implemented in EuPathDB Galaxy (Aurreocoechea et al. 2017), CAZyme annotation via HMMER/Diamond/Hotpop as implemented in dbCAN2 (Yin et al. 2012; Zhang et al. 2018), and transposable element identification via local nhmmscan v. 3.2.1 (June 2018) searches of Dfam 3.0 (Storer et al. 2021) using RepeatMasker 4.0.9 (Smit et al. 2013).

### Comparative analysis of Ext-bPI-PLCs

A long-read Nanopore assembly of *C. fimbriata* (Fourie et al. 2020) was downloaded and screened for Ext-bPI-PLCs using HMMER and SignalP as in *C. lukuohia*. The Ext-bPI-PLCs were aligned with those from *C. lukuohia* using MUSCLE in Geneious, then hand polished and trimmed to the 667 DNA characters that comprised the conserved X-domain region (Roberts et al. 2018), and the alignment used to infer a Neighbor-Joining tree in Geneious (Tamura-Nei distance, midpoint rooting). Poorly aligned Ext-bPI-PLCs with incomplete X domains (for example, due to early-terminating stop codons or frame shifts) were excluded, for a final dataset of 142 Ext-bPI-PLCs (75 from *C. lukuohia*, 66 from *C. fimbriata*, and a paralog from *Chalaropsis thielavioides* as outgroup). To compare Ext-bPI-PLC presence and organization between the two species, gene annotations of the respective Ext-bPI-PLC clusters were reciprocally transferred between assemblies using a 95% nucleotide similarity threshold with manual inspection to resolve the few ambiguous matches.

### Ext-bPI-PLC Annotation Database

Of the 49 new and 32 publicly available Microascales/Ceratocystidaceae genomes analyzed in this study (Table 1), 46 were selected to represent the diversity of Ext-bPI-PLC genes discovered in the Pfam 32.0 searches. There were 951 Ext-bPI-PLC genes identified among the 46 genomes, but 29 appeared truncated and were excluded. One Ext-bPI-PLC was identified in *Huntiella*, and homologs were identified in genomes of the three other *Huntiella* species, which were added to the database. The final “PLC annotation dataset” included 925 DNA sequences of 828 to 1,600 characters, from the start codon of the signal peptide to the stop codon. This dataset was used to annotate Ext-bPI-PLC loci across all 81 genomes with a 70% nucleotide identity threshold using Geneious (Table 1). A similar second annotation dataset was created using the six flanking genes on either side of the SC8 cluster in *C. lukuohia*, which was used to annotate the 81 genomes using a 50% identity threshold.

### Four-Gene Phylogeny of the Ceratocystidaceae

A concatenated alignment of *tef*1-α (translation elongation factor EF-1 alpha), *tub* (beta-tubulin), *mcm7* (minichromosome maintenance protein 7), and *rpl*1 (60S ribosomal protein L10a) was assembled from a family-wide alignment (Mayers et al. 2020), supplemented with orthologs extracted from genomes via local tBLASTx. Introns were removed before concatenation because of ambiguous alignment. This 92-taxon, 5975-character alignment was partitioned by gene and codon position with models as suggested by PartitionFinder 2 (Lanfear et al. 2012, 2017) powered by PhyML (Guindon et al. 2010), then used for Bayesian analysis in MrBayes 3.2.2 x64 (Ronquist et al. 2012) using a single MCMC run (one cold chain, three heated chains, 1,000,000 generations), which was sufficient for average standard deviation of split frequencies below 0.01. The final consensus tree (burnin 150,000) was visualized in FigTree v1.4.4 and manually rooted to *Fusarium graminearum*.

### Large scale bPI-PLC phylogeny

Amino acid translations of Ceratocystidaceae bPI-PLC genes (Ext-, Cyt-, and Vac-bPI-PLCs) were aligned with PI-PLC X domain alignments in the NCBI Conserved Domain Database (PF00388, Wang et al. 2023) and bacteria-like PLC (EC 4.6.1.13) sequences in InterPro (IPR000909) and anchored to the NCBI reference alignment. Using this initial set of 10,000+ sequences, preliminary neighbor-joining trees were used to select 1,010 representative sequences of clades most closely related to Ceratocystidaceae bPI-PLCs and of each major clade of bacterial bPI-PLCs (Iwasaki et al. 1998). Eukaryotic-type PI-PLCs (from NCBI and our genomes) were excluded.

Nucleotide sequences of the 1,010 proteins were extracted from the genome of origin using tBLASTx, translation-aligned, and masked upstream at the universally-conserved “IPG” of four residues (upstream from the first X domain active site, His32 in *B. cereus*) and downstream of the same 3′ residue as the *C. lukuohia*-*C. fimbriata* Ext-bPI-PLC alignment. The 597-nucleotide alignment was used to generate a neighbor-joining tree with *Listeria monocytogenes* gene AY512413 as outgroup. Branch support values were calculated in PhyML 3.0 (Guindon et al. 2010), with SMS model selection (Lefort et al. 2017) and SH-like aLRT. Major lineages were collapsed in FigTree.

### Circle tree of *Ceratocystis* Ext-bPI-PLCs

To determine whether *Ceratocystis* Ext-bPI-PLCs represent a single or multiple origins, nucleotide sequences spanning the signal peptide start codon through the end of the X domain were aligned for all available Ceratocystidaceae Ext-bPI-PLCs. This included the 925 sequences from the annotation database (except the truncated *C. lukuohia* SC10.2), plus two discovered SC2.2 sequences of North American *Ceratocystis* spp. that were not in the database, plus seven additional Ext-PI-PLCs from *Thielaviopsis*, *Endoconidiophora*, and *Davidsoniella*, for a final alignment of 933 sequences of 771 characters. A Bayesian analysis of the alignment (with the model suggested by PartitionFinder) was performed in MrBayes for 26,000,000 generations. The resulting tree was manually rooted so that *Thielaviopsis*, *Endoconidiophora*, and *Davidsoniella* sequences formed a monophyletic outgroup, which was hidden in the final tree. The tree was displayed in a circular form with tips color-coded by geographical clade of *Ceratocystis* species (North American, African, Asian-Australian, and Latin American) using the R package ggtree (Yu et al. 2017).

### Motif discovery

Putative regulatory motifs upstream of Ext-bPI-PLCs and putative cluster genes were identified using MEME v5.5.8 (Bailey and Elkan 1994; Bailey et al. 2015) in discriminative mode (ZOOPS, 6–60 nt motifs). For each gene of interest, primary sequences comprising the 1 kb immediately upstream of the start codon were compared against a control dataset of all non-cluster genes (*C. lukuohia*) or a random selection of 10,000 estimated ORFs in the genome (*B. basicola* and *Ch. thielavioides*). Analyses focused on the six conserved cluster flanking genes (a, b, c, h, i, j), cluster Ext-bPI-PLCs genes, and other genes. Other searches tested alternative sets, including the non-cluster Ext-bPI-PLCs in *B. basicola* and *C. lukuohia*. Significance thresholds were evaluated using sequence-shuffled runs.

The identified enriched motifs were searched for putative ontology against the *Schizosaccharomyces pombe* GOMo database (Buske et al. 2010) and compared to known fungal TF binding motifs in JASPAR “NON-REDUNDANT CORE Fungi 2024” (Rauluseviciute et al. 2024) using Tomtom (Gupta et al. 2007). Genome-wide motif occurrences were mapped with FIMO (Grant et al. 2011) and pruned by q-value (typical threshold 0.01).

## RESULTS

### Long and short read assemblies of *C. lukuohia*

Long-read assemblies (HGAP4 and Canu) of *C. lukuohia* C4212 were substantially improved over the short-read SPAdes assembly and the hybrid (SPAdes) assembly of the same isolate (Supplemental Table 1). The Canu assembly had 84 unitigs excluding mitochondrial (low GC) unitigs, 33 nuclear contigs, N50 = 2.9 Mb, and an estimated total nuclear genome of 32.1 Mb. Sixteen of the contigs had unique telomere/subtelomeric regions (Supplemental Table 1). Short-read mapping (Geneious “map to reference”) supported each of the 16 telomeres, suggesting at least eight chromosomes in the genome.

A curated set of 12 supercontigs (Fig. 1; Supplemental Table 2) was assembled by linking 84 Canu unitigs using the HGAP and short-read SPAdes contigs. This assembly (JABSNW000000000) was wholly supported by paired-end read mapping, except for 3800 bp of ambiguity at the 3′ end of supercontig 6, which shared homology with the 3′ region of SC10 and the middle of SC1. Six supercontigs had telomeres at both ends, suggesting they represent full chromosomes (Fig. 1; Supplemental Table 2). Supercontig 9 (1,423 kb, telomere-to-telomere) likely represents the smallest whole chromosome, with the very small (28–608 kb) supercontigs 10, 11 and 12 likely representing misassembled fragments. Telomere distribution, repeat features and gene-model summaries for each supercontig (Supplementary Tables 2 and 3) suggest that at least the nine largest supercontigs represent typical nuclear chromosomes, whole or partial, consistent with previous estimates of seven to nine linkage groups or chromosomes in *Ceratocystis* (Fourie et al. 2019; Fernandes et al. 2022; Nakamoto et al. 2026). Putative transposable elements (TEs) were widespread among the supercontigs (Fig. 1), often in clusters, occasionally near the end of a supercontig, where they may have impeded unitig extension.

**Figure 1.**
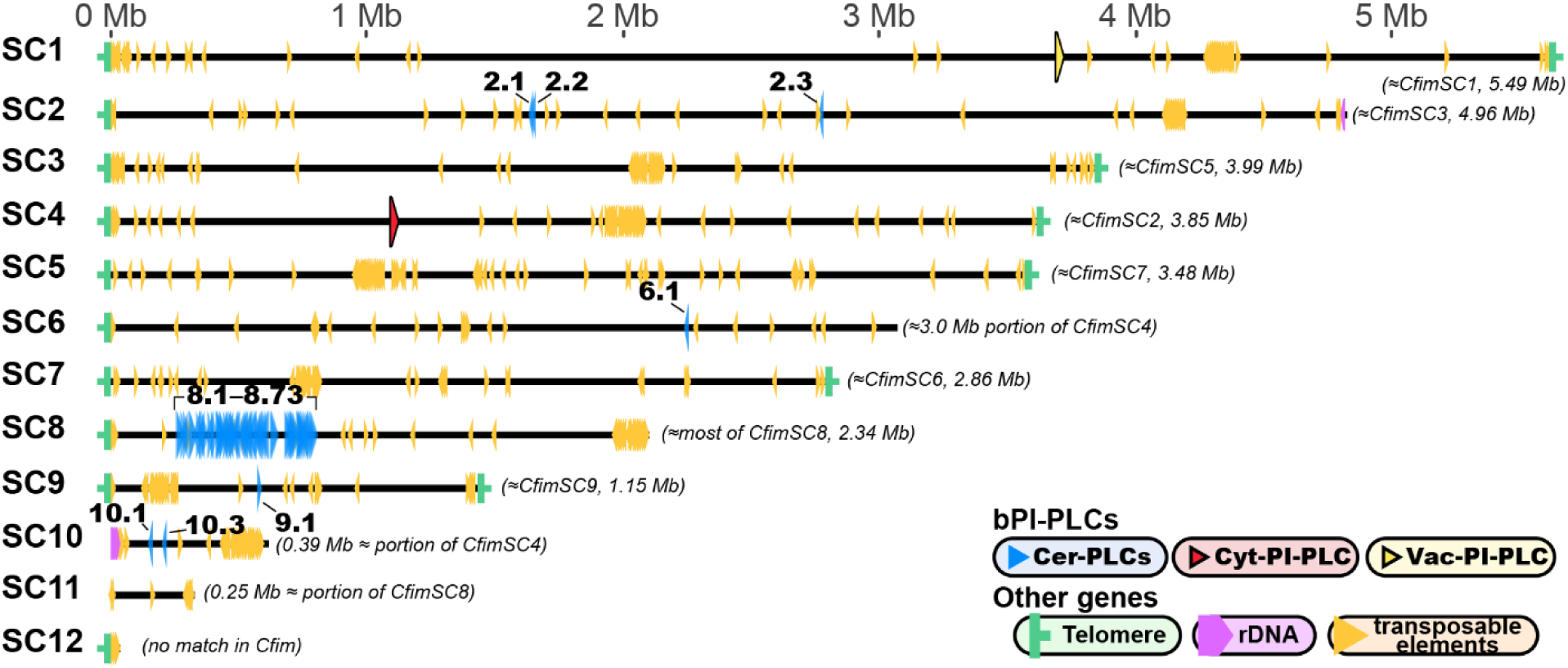
Distribution of putative PI-PLC genes, transposable elements, rDNA operon genes, and telomeres of *Ceratocystis lukuohia* C4212 (CBS 142792) on the 12-supercontig assembly (JABSNW000000000). Bacterial-phosphatidylinositol-specific-phospholipase-C-like genes (EC 4.6.1.13; bPI-PLCs) with signal peptides (Cer-PLCs) are indicated by blue triangles and numbered as in Supplemental Table 4. The “Cer-PLC cluster” (Cer-PLC genes 8.1–8.73) is evident on supercontig 8. The “cytoplasmic” Cyt-PI-PLC and “vacuolar” Vac-PI-PLC genes are indicated in red and yellow, respectively. Equivalent supercontigs in *C. fimbriata* are given, along with their length. Scale bar is megabases (Mb).

Supercontig 10 had a particularly high number of TEs, again suggesting that it may be a misassembled contig. Two complete copies of the rDNA operon at the beginning of supercontig 10 and a fragment of the 28S rDNA gene at the end of supercontig 2 indicate that the very long rDNA repeat region (Lofgren et al. 2019) was not fully assembled. Overall, 7,556 genes (AUGUSTUS) comprising 6,195 orthologs (OrthoMCL) achieved an odb9 BUSCO completeness of 89.7%. Of the 7,556 genes, 10% were predicted to have signal peptides, with supercontigs 8 and 9 particularly enriched for signal peptides (Fig. 1, Supplemental Tables 2 and 3). The distribution of total predicted and carbohydrate active (CAZyme) genes (Supplemental Table 3) suggests that the nine largest supercontigs do not represent dispensable chromosomes (DE). The eight largest supercontigs were larger than typical fungal DEs, which are generally less than 2 Mb (Mehrabi et al. 2017).

### Identification of PI-PLCs among the 12 *C. lukuohia* supercontigs

A genome-wide HMMER search for PI-PLC-X domain proteins (PF00388.19) identified 88 PI-PLC candidates among the translated ORFs of the supercontigs after combining two overlapping ORFs and excluding two ORFs for which putative translations had no significant BLASTn/BLASTp hits. A separate AUGUSTUS/BLASTp (nr database) search predicted 87 PI-PLCs, including one not found by HMMER. Cross-checking between the HMMER and AUGUSTUS search results identified 89 PI-PLCs and helped refine start/stop boundaries for 14 of the genes. The 89 candidates were numbered by supercontig and position, e.g., PI-PLC 8.2 for the second PI-PLC-X gene on SC8 (Supplemental Tables 4 and 5; Figs. 1 and 2).

**Figure 2.**
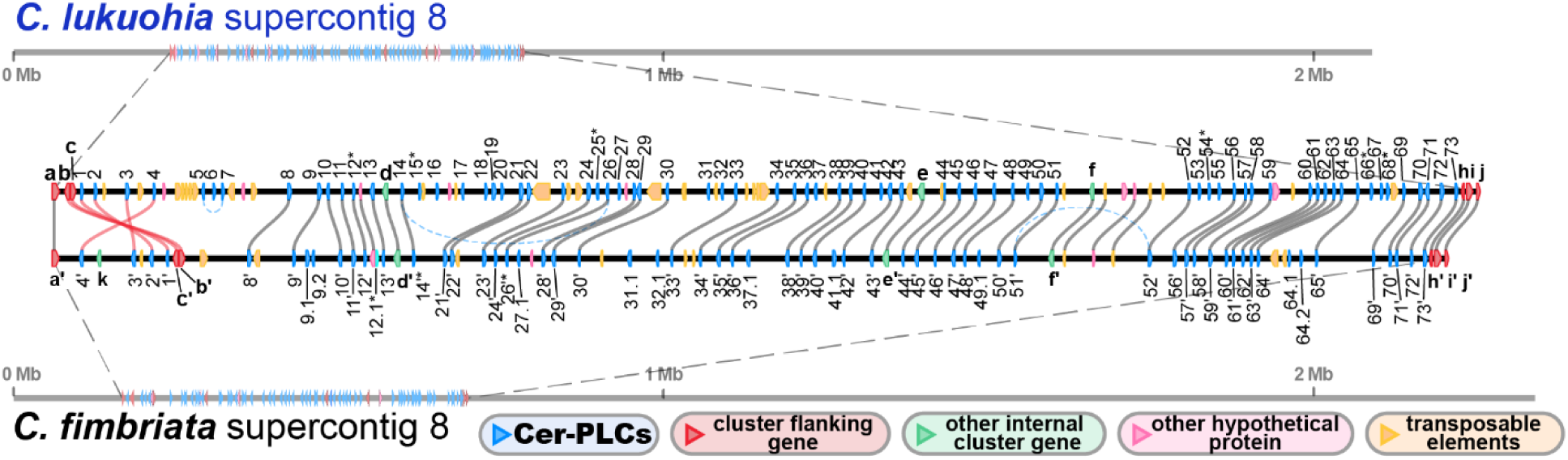
Detailed maps and comparative organizations of genes in the Cer-PLC clusters of *Ceratocystis lukuohia* (top) and *C. fimbriata* (bottom). Both clusters are connected to their respective positions on their full supercontigs of origin (supercontig 8 in both species) by dotted lines at their margins. Genes are color-coded as per legend. Cer-PLCs are numbered according to Supplemental Tables 4 and 5; partial Cer-PLC genes are marked with an asterisk (*). Selected non-PLC genes (cluster flanking and internal cluster genes) are labelled with letters according to Table 6 and Figure 3. Closely orthologous genes (≥95% similarity) of the two species are connected by gray lines if in the same order and orientation, red lines if part of an inversion, or dotted blue lines if within the species. *C. lukuohia* assembly JABSNW000000000 (isolate C4212/CBS 142972); *C. fimbriata* assembly from Fourie et al. (2020) (isolate CMW 14799/CBS 114723). Scale bar in megabases (Mb).

Six of the 89 genes coded for the PI-PLC-X domain but were excluded from further study because their products were identified as eukaryotic PI-PLCs (EC 3.1.4.11; hereafter “ePI-PLC”) or as other, unrelated eukaryotic enzymes (Supplemental Table 4). The remaining 83 candidates were classified as prokaryotic PI-PLCs (EC 4.6.1.13; hereafter “bPI-PLC”) based on BLASTp searches and the KEGG database (BlastKOALA v. 2.2, Kanehisa et al. 2016). Two of the 83 genes appeared to be single-copy genes found in related ascomycetes; one was cytoplasmic (Cyt-bPI-PLC) and the other lysosomal/vacuolar (Vac-bPI-PLC) based on Deeploc searches. The remaining 81 represented a clade of unique paralogs (hereafter “Cer-PLCs”), distinguished from the other Ext-PI-PLC genes found in some other genera of the family. Translations of 74 of the 81 genes had complete X domains, N-terminal Sec/SPI signal peptides (Almagro Armenteros et al. 2019), and extracellular localization by Deeploc, whereas four (8.12, 8.15, 8.25, and 8.68) were N-truncated without functional signal peptides and three (8.54, 8.66 and 10.2) were C-truncated within the X domain (Supplemental Table 4). Three of the 74 full Cer-PLCs (Supplemental Table 4) showed high sequence identity with three of the four PLC proteins detected in the secretome of *C. cacaofunesta* (Molano et al. 2018).

### The Cer-PLC cluster in *C. lukuohia*

Cer-PLCs were highly concentrated on supercontig 8 (Figs. 1, 2 and 3), where 73 of the 81 full or partial Cer-PLCs (90%) occur within a single 524-kb region. Besides the 73 Cer-PLCs, 48 cluster genes were predicted by AUGUSTUS, mostly identified as transposable elements or hypothetical proteins. Five internal genes, as well as three 5’ and three 3’ genes immediately flanking the cluster of Cer-PLCs, were tentatively identified by BLASTp searches of the nr database and were assigned shorthand letters from ‘a’ to ‘j’ (Figs. 2 and 3; Supplemental Table 6). Most notable of these was ‘a’, coding for a putative CeGAL Zn2-C6 transcription factor (Mayer et al. 2023).

**Figure 3.**
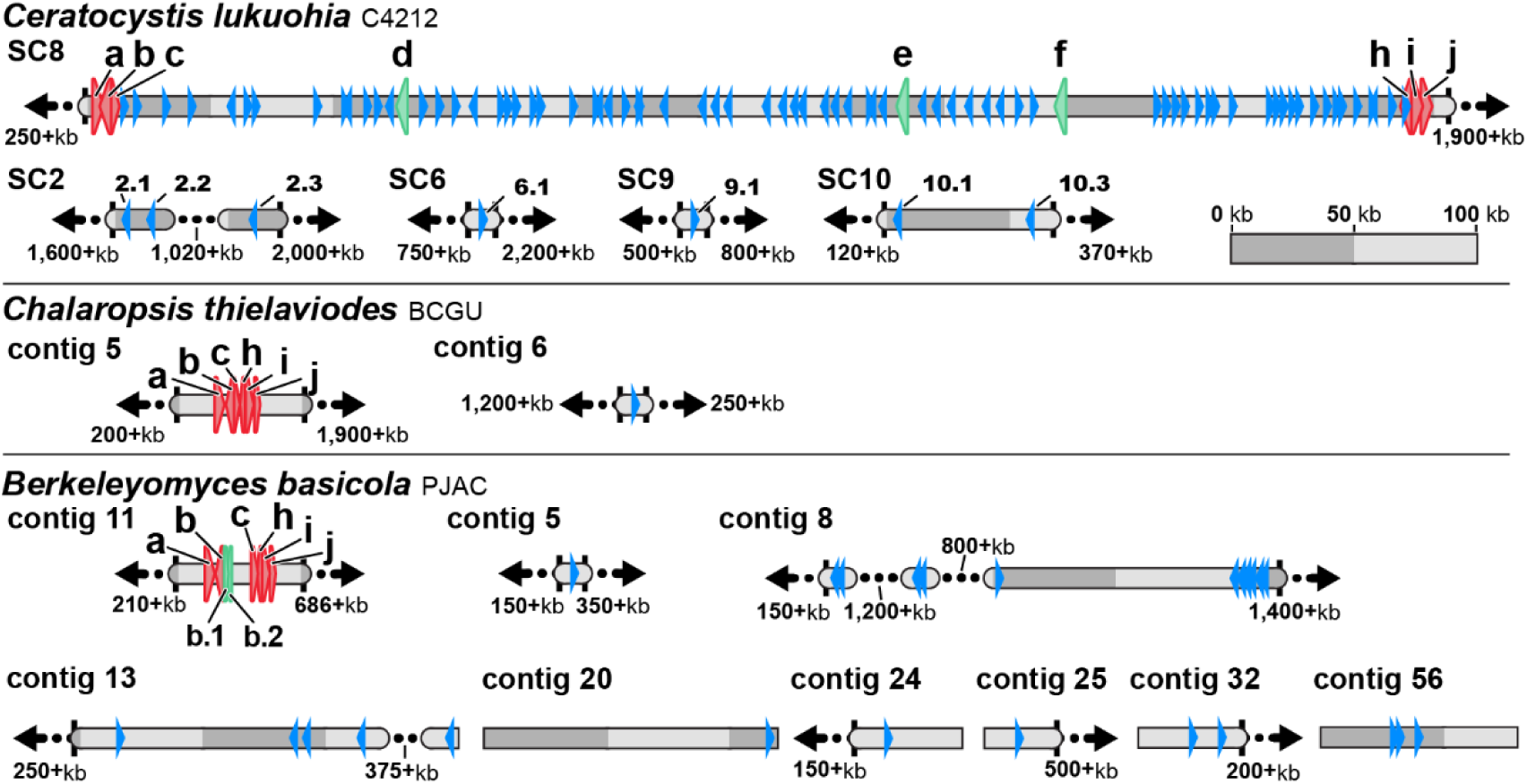
Schematic depictions of the locations of identified Cer-PLCs (blue) in the genomes of *Chalaropsis thielavioides, Ceratocystis lukuohia*, and *Berkeleyomyces basicola*, as well as the Cer-PLC cluster in *Ceratocystis* and the homologous cluster site in *Chalaropsis* and *Berkeleyomyces*. In *Chalaropsis*, only a single Cer-PLC was identified, and six genes (a, CeGAL zinc cluster transcription factor; b, AAA+ ATPase; c, a hypothetical protein; h, rab-like small GTPase; i, ATP synthase regulation protein nca2; and j, fatty acid hydroxylase) exist as an uninterrupted cluster. In *Ceratocystis*, the vast majority of Cer-PLCs are in the SC8 Cer-PLC cluster, which is flanked by a, b, and c on its 5′ end and by h, i, and j on its 3′ end. *Berkeleyomyces* has fewer PI-PLCs than *Ceratocystis*, and they are spread more widely across its contigs. The *Berkeleyomyces* a,b,c,h,i,j cluster has two additional identified genes inserted between b and c: b.1, a metallo-beta-lactamase type-2 like family gene, and b.2, an NMRA-like family protein. The flanking genes (a,b,c,h,i,j) are in red, and other non-Cer-PLC genes are in green. All contigs are to scale (as per key, in kilobases), though dotted lines or arrows represent portions excluded from some contigs in this illustration.

The five Cer-PLCs found on SC6, SC9, and SC10 (Figs. 1 and 3) have signal peptides and standard bPI-PLC domains similar to the Cer-PLCs in the SC8 cluster, though the SC10.2 Cer-PLC was truncated (Supplemental Table 4). The three Cer-PLCs on SC2 were distinct (Fig. 4B): SC2.1 and SC2.2 were near each other, and their translations each had extensions ending in a proline-rich intrinsically disordered region (per InterProScan). The translation of SC2.3 was nearly twice the length of typical Cer-PLCs due to the addition of a C-terminal metallo-hydrolase/oxidoreductase domain (DUF4336 family, per InterProScan).

**Figure 4.**
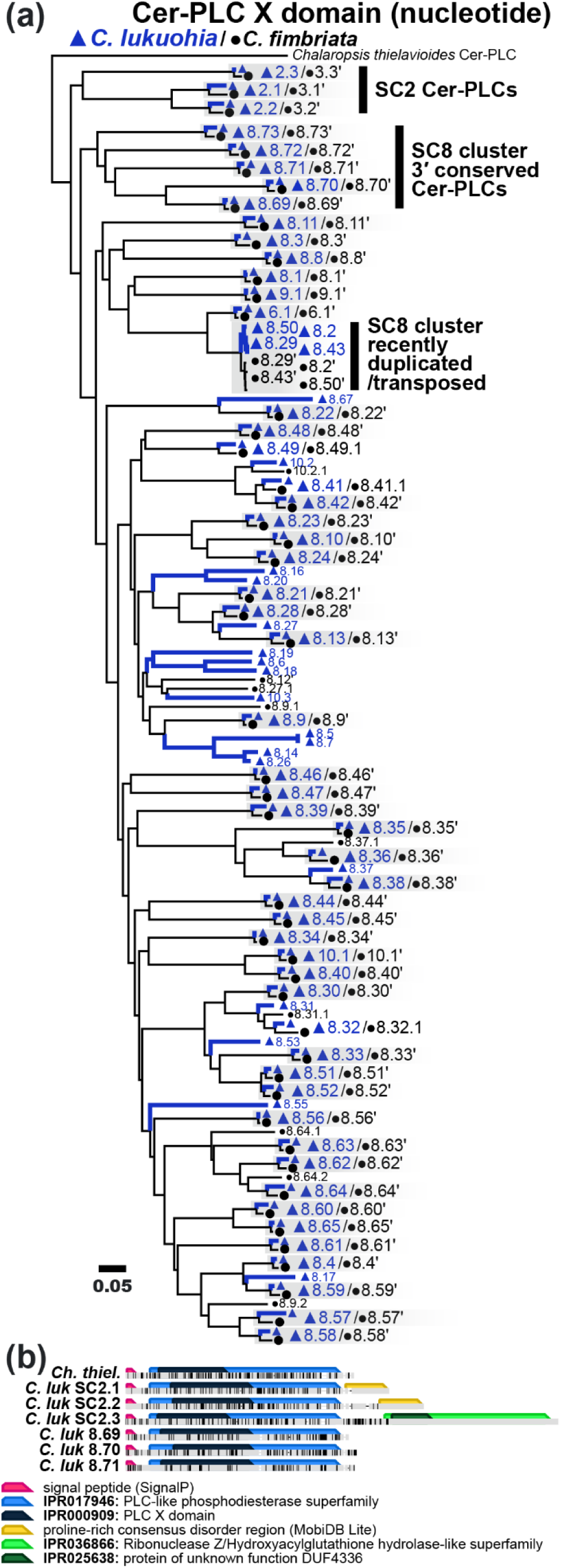
Neighbor-Joining nucleotide tree of the PLC-X domain of 141 Cer-PLC genes from *Ceratocystis lukuohia* (blue triangles and branches) and *C. fimbriata* (black circles and branches). (A) The single Cer-PLC found in *Chalaropsis thielavioides* is used as the outgroup. Cer-PLC genes are labelled as in Supplemental Tables 4 and 5, where numbers left of decimal points indicate supercontig and numbers right of decimal points indicate relative gene positions. Gray boxes highlight closely orthologous Cer-PLCs (≥95% similarity). Bar represents 0.05 substitutions/site. (B) Comparison of the SC2-type Cer-PLCs with C-terminal extensions to other Cer-PLCs.

### Comparison of *C. fimbriata* and *C. lukuohia* PI-PLCs and the Cer-PLC cluster

The nine supercontigs of the *C. fimbriata* assembly (Fourie et al. 2020) appeared homologous to the nine largest supercontigs of *C. lukuohia*, and the distribution of their bPI-PLCs were similar (Supplemental Tables 4 and 5). Screening the 178,357 translated ORFs of *C. fimbriata* yielded 80 candidates with PI-PLC X domains, but two were excluded due to lack of functional homologs or incomplete X domains, leaving 78 candidate PI-PLCs. Five of these were ePI-PLCs, two were homologs of *C. lukuohia* Cyt- and Vac-bPI-PLCs, and three encoded unrelated genes (Supplemental Table 5). The remaining 69 represent Cer-PLCs, which were each named after their orthologs in *C. lukuohia* (> 95% identity) plus a prime symbol, or by decimal names relative to their positions on the *C. lukuohia* supercontigs (e.g., 8.12.1 in *C. fimbriata* is between the positions for 8.12 and 8.13 in *C. lukuohia*) (Fig. 2, Supplemental Table 5). Four *C. fimbriata* Cer-PLCs were truncated, leaving 65 full-length Cer-PLCs (Supplemental Table 5), each encoding a signal peptide and with extracellular localization.

The Cer-PLC cluster is on the eighth largest supercontig and in roughly the same position in each species according to MAUVE alignment. Fifty-eight of the 65 *C. fimbriata* Cer-PLCs clustered on CfimSC8 (Fig. 2). The two long-read assemblies of *C. fimbriata* (Fourie et al. 2020, Stahr et al. 2024) had CfimSC8 lengths of 2.34 Mb and 2.33 Mb, respectively, but telomeres were not identified in these assemblies. The *C. lukuohia* SC8 was smaller (2.09 Mb), likely due to a high density of TEs at the 3′ end, which showed homology to the short and poorly assembled SC11 (Fig. 1). The cluster (including flanking genes) is slightly smaller in *C. fimbriata* (530,449 bp) than in *C. lukuohia* (542,833 bp). The orthologous pairs of Cer-PLCs (>95% similarity) showed a broad similarity in gene ordering within the cluster, but there were multiple insertions, deletions, and a putative inversion involving groups of Cer-PLCs and their non-coding intergenic spaces (Fig. 2).

Besides TEs and Cer-PLC genes, the genes flanking the cluster and the five identified internal genes of the two species largely shared orthology and gene order, with the notable exception of a large, inverted region (36,648bp in *C. lukuohia* and 41,999 bp in *C. fimbriata*) after the transcription factor ‘a’. The other two 5’ flanking genes (b and c) and four orthologous Cer-PLCs were involved in the apparent inversion (Fig. 2), along with the addition of gene ‘k’ in *C. fimbriata,* which encodes a HET-E1 vegetative incompatibility protein similar to *Podospora anserina* HET-E1 (Q00808, Daskalov et al. 2023; Espagne et al. 2002). Other HET-E1 genes exist in the nuclear genomes of *C. fimbriata* and *C. lukuohia*, but none were a close match for ‘k’. Other internal cluster genes and 3’ flanking genes (d, e, f, h, i, j) were found in both species (Fig. 2, Supplemental Table 6)).

### Cer-PLC gene diversity within *C. lukuohia* and *C. fimbriata*

The full Cer-PLCs of *C. lukuohia* (74 genes) and *C. fimbriata* (67 genes) aligned unambiguously from the start of the signal peptide through the X domain. However, each Cer-PLC had a unique DNA sequence, and only two (8.5 and 8.7 in *C. lukuohia*) encoded identical amino acid sequences. The neighbor-joining tree of the 141 DNA sequences (Fig. 4) demonstrated one-to-one correspondence between most Cer-PLCs of the two species, with 58 orthologous pairs (>97% identity) that preserved the relative order within the clusters, except for the putative 5’ inversion (Fig. 2; Supplemental Tables 4 and 5). Fifteen *C. lukuohia* Cer-PLCs had no clear counterpart in *C fimbriata*, but two of those (8.14 and 8.26) matched truncated Cer-PLCs in *C. fimbriata* (8.14’ and 8.26’). Seven *C. fimbriata* Cer-PLCs had no equivalent in *C. lukuohia,* but one (8.12’) matched the truncated 8.12 (Fig. 2).

The NJ tree of the 141 Cer-PLCs (Fig. 4) showed that the three earliest-diverging ortholog pairs are the unique Cer-PLCs found on SC2 in *C. lukuohia* and SC3 of *C. fimbriata*, with the translations of the *C. fimbriata* genes showing the same C-terminal extensions as the comparable genes in *C. lukuohia*. The other Cer-PLCs form a monophyletic group of cluster Cer-PLCs as well as a few Cer-PLCs on other respective supercontigs (Fig. 4). Within the clade comprising the cluster Cer-PLCs, five ortholog pairs (8.69–8.73) form an early-diverging subclade, and the five pairs are located at the 3’ end of the respective clusters (Figs. 2 and 4). Four nearly-identical (recently diverged) Cer-PLCs in *C. lukuohia* (SC8.2, SC8.29, SC8.43 and SC8.50) are distributed across the SC8 cluster (Fig. 2), and this group is sister to the orthologous group of four nearly-identical Cer-PLCs in *C. fimbriata* (Figs. 2 and 4; Supplemental Tables 4 and 5).

### Genome Assemblies of Ceratocystidaceae and Phylogeny

Assemblies with fewer contigs tended to have larger estimated genome sizes, as expected. For example, SPAdes assemblies of *C. lukuohia* were estimated to be 30.6–30.7 Mb (7429 –7480 estimated Augustus genes), whereas the long-read assembly of C4212 totaled 31.7 Mb and 7407 estimated genes (Table 1). The same pattern was evident for *C. fimbriata* and *B. basicola* assemblies (Table 1). The long-read genome assembly of the AAC species *C. huliohia* (Nakaimoto et al. 2026) had 7006 estimated genes across 29.3 Mb, whereas our five short read-assemblies of *C. huliohia* had an estimated 7017–7060 Augustus genes across only 27.9 Mb.

The inferred phylogeny of the Ceratocystidaceae using four gene sequences extracted from each genome (Fig. 5) showed expected evolutionary relationships within the family (de Beer et al. 2014, 2017; Nel et al. 2018; Mayers et al. 2020). The monophyly of the *Berkeleyomyces-Chalaropsis-Ceratocystis* (*Berk-Chal-Cer*) Clade was strongly supported, as was separation of the four geographic clades within *Ceratocystis* (Johnson et al. 2005; Mbenoun et al. 2014; Li et al. 2017; Liu et al. 2018; Holland et al. 2019; Harrington et al. 2024). The *Thielaviopsis*-*Endoconidiophora*-*Davidsoniella* (*Thiel-Endo-David*) Clade formed a sister group to the *Berk-Chal-Cer* Clade (Fig. 5a).

**Figure 5.**
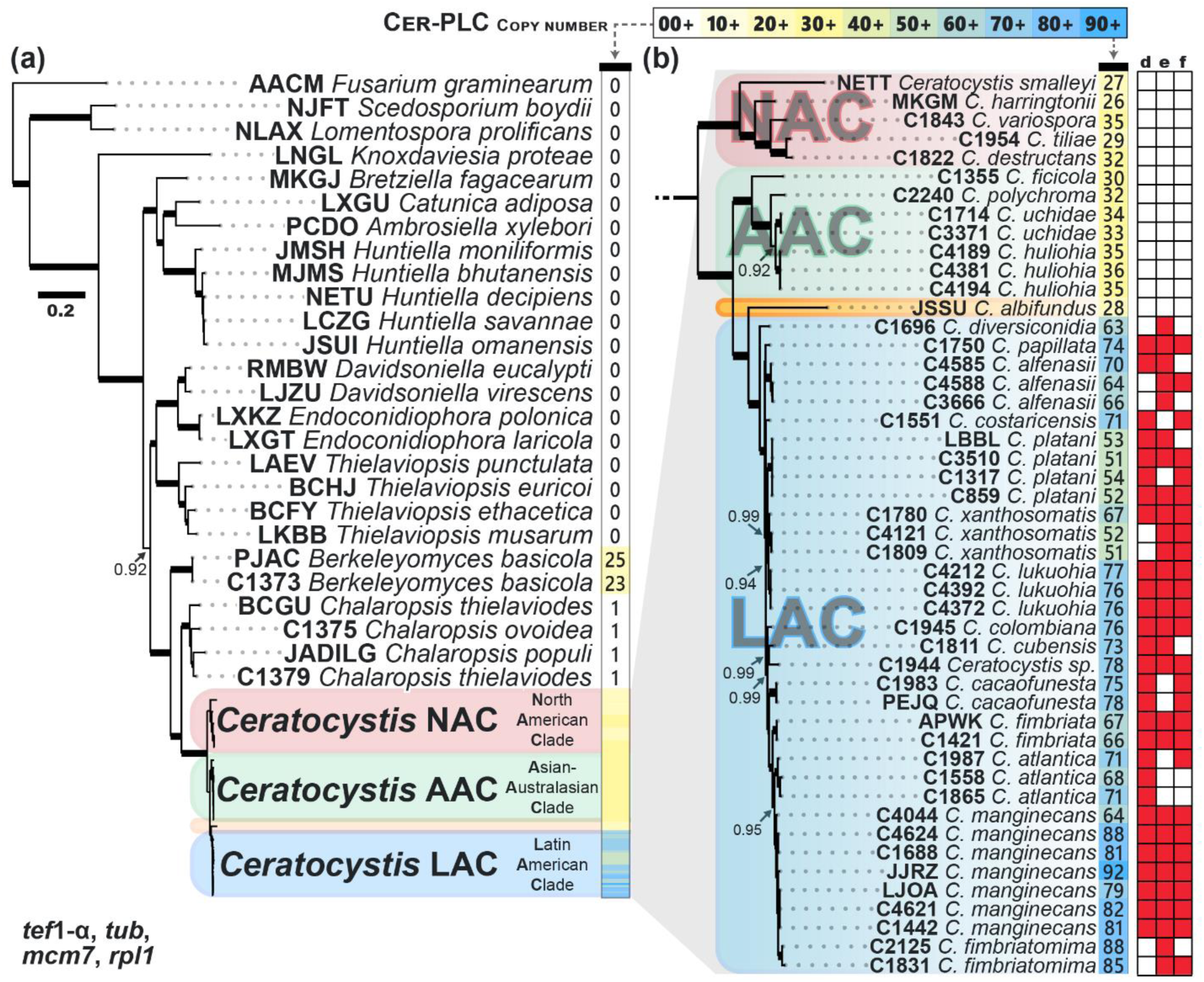
Bayesian four-gene phylogeny (*tef*1-α, *tub*, *mcm7*, and *rpl1*) and estimated number of Ext-PI-PLCs genes or Cer-PLC genes in each genome based on searches with an annotation database. (A) The Ceratocystidaceae family. (B) Expanded subset of A comprising the genus *Ceratocystis* and its geographic subclades: North American Clade (NAC, in red), Asian-Australasian Clade (AAC, in green), and African lineage (*C. albifundus*, in orange) and Latin American Clade (LAC in blue). Numbers to the right of each species represent the number of Cer-PLC genes we identified in its genome, color-coded from yellow (lowest) to green (intermediate) to blue (highest), per the scale at top. The three columns at right indicate the presence (red) or absence (white) of three internal genes within the Cer-PLC cluster (d, e, and f, per Supplemental Table 6). Posterior probability supports ≥0.90 are indicated; thick branches indicate values of 1.0. Scale bar = 0.2 estimated substitutions per site.

*Chalaropsis* has the smallest genomes (23.2–23.9 Mb and 6394–6878 genes) among the taxa studied, smaller than the 25.5 Mb and 7025 genes estimated for *B. basicola* (Table 1). In *Ceratocystis,* the Latin American Clade (LAC) had the largest and most variable genome sizes (28.8–33.2 Mb, 7092–7966 genes). The North American Clade (NAC) had genomes of 25.8–27.5 Mb and 6620–6994 genes. The Asian-Australasian Clade (AAC) had genomes of 26.7–27.9 Mb and 6840–7781 genes, except for the large estimate of 29.7 Mb for the recent *C. huliohia* assembly (Nakimoto et al. 2026). The African species *C. albifundus* had a genome of 26.9 Mb and 6689 genes.

### Identification of Ext-bPI-PLCs in Ceratocystidaceae

The HMMER screening of predicted ORFs identified a single Ext-bPLC in each assembly of *Endoconidiophora*, *Davidsoniella*, and *Thielaviopsis*, except for *T. punctulata,* which had none by this method (Table X). Only one *Huntiella* (*H. bhutanensis*) had a detected Ext-bPI-PLC by this method. There were 21 Ext-bPI-PLCs identified in *Berkeleyomyces basicola* and a single Ext-bPI-PLC in each *Chalaropsis* assembly (Table 1). A range of 37 to 102 bPI-PLC X-domain candidates were detected in each of 21 *Ceratocystis* genomes, and 24 to 77 had signal peptides (Ext-bPI-PLCs). These Ext-bPI-PLC estimates (Table 1) were highest in LAC species (46–77 Ext-bPI-PLCs) versus species in the other clades (AAC = 24–33; NAC = 25–30; *C. albifundus* = 27). By manual alignment of amino acid sequences, each of the *Berk-Chal-Cer* Ext-PI-PLCs was identified as a Cer-PLC. No Ext-PI-PLC was detected in *Ambrosiella*, *Bretziella*, *Catunica*, or other Microascales (*Knoxdaviesia*, *Scedosporium* and *Lomentospora*), nor in *Fusarium graminearum* (Table 1).

Domain-based ORF screening can miss fragmented or poorly predicted ORFs, so a more thorough search for Ext-bPI-PLCs was conducted against 83 genomes using an Ext-bPI-PLC/Cer-PLC annotation database of 925 sequences that represented the diversity of Ext-bPI-PLCs identified in the family through HMMR searches. Annotations at the relatively permissive 70% identity generally identified more Ext-bPI-PLCs than did the HMMER searches, especially for *Ceratocystis* genomes (Table 1). A single Ext-bPLC was detected in *Endoconidiophora*, *Davidsoniella*, *Thielaviopsis* (including *T. punctulata*) and the four species of *Huntiella* (Fig. 5A). The annotation database identified 23–25 Cer-PLCs in the *B. basicola* assemblies and a single copy in the *Chalaropsis* assemblies.

The annotations gave consistent results across the different assemblies of the same *Ceratocystis* isolate (Table 1). For example, only 55 Cer-PLCs were identified via HMMER in the short-read *C. fimbriata* genome (APWK, Wilken et al. 2013), versus 67 of the same assembly with the annotation database, 66 annotations for our own assembly of the same isolate (C1421), and 66 annotations for the long-read assembly (Fourie et al. 2020) of the same isolate (Table 1, Fig. 5B). As in the HMMR estimates, the Cer-PLC estimates based on annotation varied widely across the genus, with the widest range of counts (51–92) found in the LAC (Table 1, Fig. 5B). Within the LAC, Cer-PLC estimates of the host-restricted *C. platani* (two isolates from Europe and two from North America) were similar (51–54), whereas Cer-PLC estimates for isolates of *C. xanthosomatica* varied by host strain (51 and 52 in *Syngonium* isolates, 67 in the *Xanthosoma* isolate). The largest Cer-PLC counts and intraspecies variation occurred in *C. manginecans*: an indigenous isolate of a seedling blight strain on *Carapa* from the Amazon Basin (Valdetaro et al. 2019) had only 64 Cer-PLCs, whereas there were 92 Cer-PLCs in the invasive and highly aggressive Omani strain (Fig. 5B).

### Comparisons of bPI-PLCs in Ascomycetes and Prokaryotes

Phylogenetic analysis of prokaryotic-type PI-PLCs (bPI-PLCs) from fungi and bacteria resolved several well-supported fungal lineages of PI-PLCs (Fig. 6). Most bPI-PLCs of bacteria clustered with the *Listeria monocytogenes* bPI-PLC, which was used as the outgroup. *Streptomyces* bPI-PLCs, which are known to be distinct from the bPI-PLCs of other bacteria (Moser et al. 1997; Iwasaki et al. 1998), formed a poorly-supported clade embedded among the bPI-PLCs from fungi. This ambiguous placement appeared to be a significant factor in the low support for the surrounding clades of fungal bPI-PLCs (Fig. 6).

**Figure 6.**
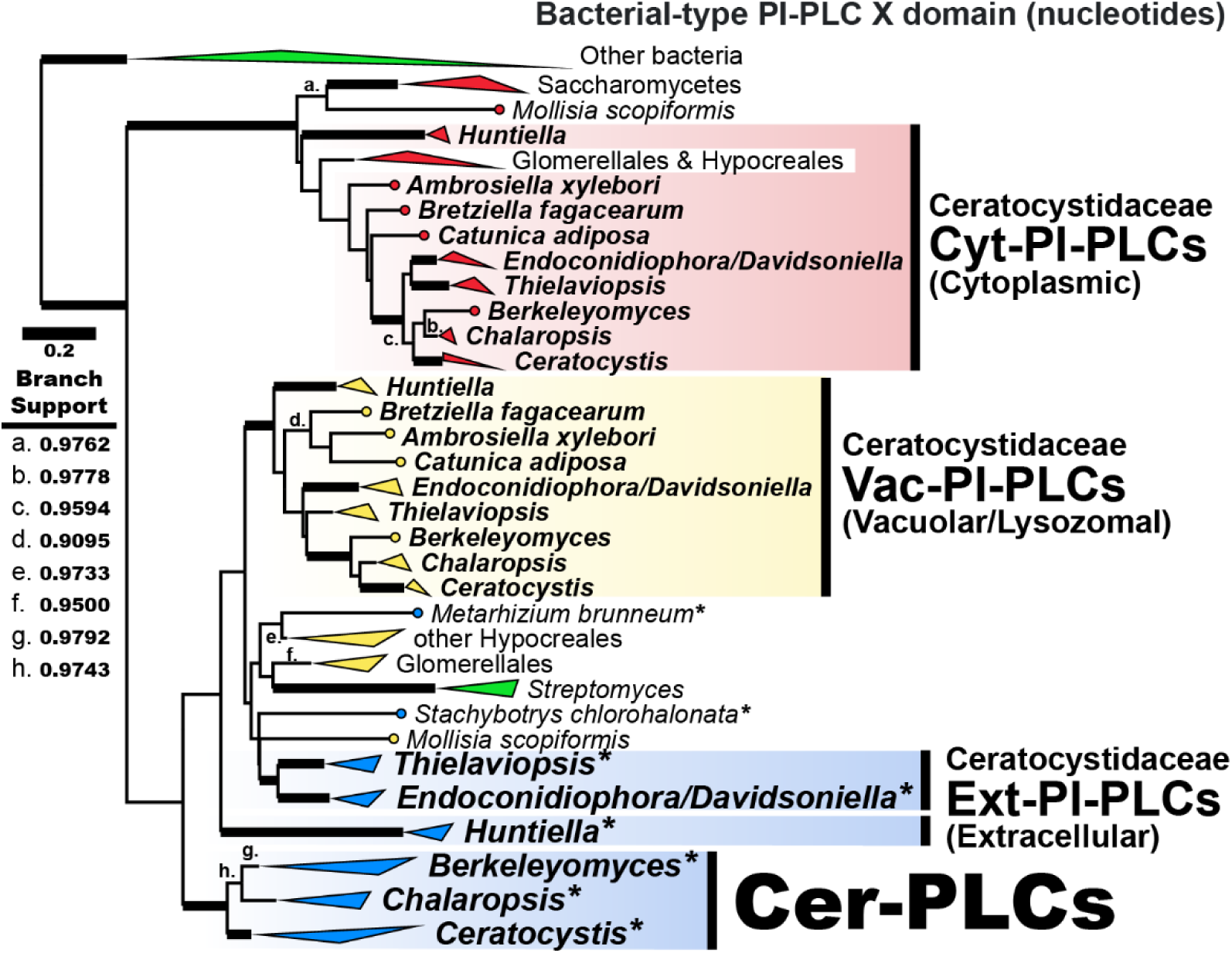
Phylogenetic diversity of bPI-PLC-like and Cer-PLC genes across fungi and bacteria. A simplified gene phylogeny from nucleotide Neighbor Joining analysis of >900 bPI-PLC-like (EC. 4.6.1.13) genes from Ceratocystidaceae, other ascomycetes, and bacteria. Triangles represent collapsed monophyletic clades: bacterial bPI-PLCs (green), fungal cytoplasmic bPI-PLCs (i.e. Cyt-PI-PLCs, red), fungal vacuolar bPI-PLCs (i.e. Vac-PI-PLCs, yellow), fungal extracellular bPI-PLCs (i.e. Ext-PI-PLCs, blue) and Cer-PLCs (also blue). Branch labels indicate branch support from PhyML analysis, as indicated in the key. Branches with more than 0.99 support are thickened. Scale: 0.2 substitutions per site. The full tree is available as Supplemental Figure 2.

Among Ceratocystidaceae bPI-PLCs, clades of two conserved, single-copy genes with different cellular locations based on DeepLoc were strongly supported: a cytoplasmic (Cyt-bPI-PLC) clade and a vacuolar/lysosomal (Vac-bPI-PLC) clade (Fig. 6). The topology of the genera and species within each of the two gene clades broadly agreed with the four-gene tree (Fig. 5). Each clade included bPI-PLCs of other ascomycetes, though the Vac-bPI-PLC clade appeared to be restricted to the Hypocreomycetidae.

Outside of Ceratocystidaceae, Ext-bPI-PLCs were only found in two species of Hypocreales (Fig. 6). There were two clades of extracellular (signal peptide) bPI-PLCs of the Ceratocystidaceae: the single-copy orthologs of *Thielaviopsis*, *Endoconidiophora*, and *Davidsoniella* and the Cer-PLCs of *Berkeleyomyces, Chalaropsis,* and *Ceratocystis* (Fig. 6). By amino acid alignment, these two clades of Ext-bPI-PLCs appeared more similar to the Vac-bPI-PLCs than to the Cyt-bPI-PLCs. However, relationships among these lineages were only weakly supported, as was the placement of the *Huntiella* Ext-bPI-PLCs (Fig. 6).

### Putative translations of Ext-PI-PLC and Cer-PLC genes

The bPI-PLC protein translations were examined for six canonical residues in the functional domain that are associated with catalytic activity and phosphatidylinositol specificity (Fig. 7). These residues were broadly conserved among the bPI-PLCs of bacteria and were present in most of the bPI-PLCs of fungi. Four of the six residues were unanimously conserved, and two nearly so (Fig. 7). In contrast, the Cer-PLCs from *Berkeleyomyces, Chalaropsis*, and *Ceratocystis* showed extensive and distinct divergences that suggest that Cer-PLCs have altered catalytic properties and/or substrate interactions in comparison to canonical bPI-PLCs (Fig. 7, Supplemental Figure 1). Using residue numbering of *Bacillus cereus*, nearly all Cer-PLCs replaced the positively-charged Arg69 with the uncharged Ser or Thr, and most substituted Lys115 with Gln. Substitutions were also common for Asp33 and His82, with few substitutions for His32, and Asp274 was conserved (Fig. 7).

**Figure 7.**
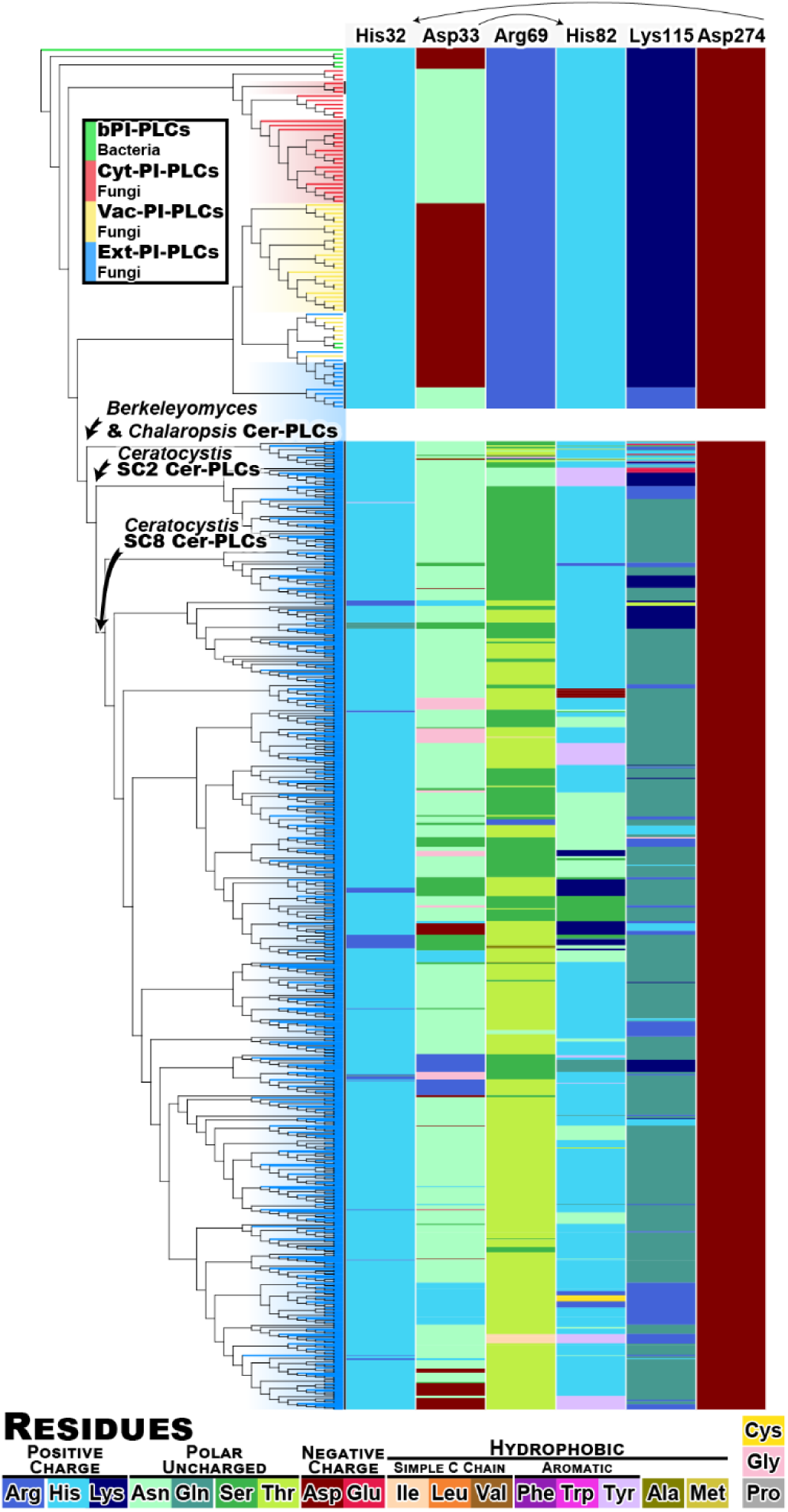
Variation in bPI-PLC and Cer-PLC critical active-site residues among *Ceratocystis*, other fungi, and bacteria. Left: gene phylogram derived from the topology of the full tree depicted in Fig. 6. Right: colored columns show conservation of six important amino acid sites for bPI-PLCs as defined in *Bacillus cereus*, with amino acid colors as per the legend. The stems of the *Ceratocystis* Cer-PLCs from supercontig 8 and supercontig 2 are indicated, as well as the stem of the Cer-PLCs from *Berkeleyomyces* and *Charlaropsis*.

Simulations by Molano et al. (2018) predicted a different set of five binding residues for their PLC s111.3 (our Cer-PLC 8.33) that could potentially accommodate inositol binding. This alternative includes Thr95 (homologous to *B. cereus* Arg69), Gln140 (*B. cereus* Lys115), and His58 (*B. cereus* His32), and these three residues are generally well conserved among Cer-PLCs (Fig. 7). The other two potential binding residues have homologs in the PI-PLC of *Listeria monocytogenes* (Molano et al. 2018) but not in the PI-PLC of *B. cereus*. Their Arg138 residue is well conserved in the Cer-PLCs of *Ceratocystis* and *Berkeleyomyces* (alignment position 157 in Supplemental Fig. 1) but not in *Chalaropsis*. Their Ser227 was not conserved in our dataset (Supplemental Figure 1, alignment position 253), further questioning if Cer-PLCs are phosphatidylinositol-specific.

Broadly conserved cleavage sites for signal peptides were seen for the Cer-PLCs. Otherwise, the signal peptides varied greatly and were less conserved than the X domain (Supplemental Figure 1), as is typical for signal peptides (Li et al. 2009).

### Cer-PLC Evolution

A phylogenetic analysis of 933 DNA sequences of representative Cer-PLCs and the Ext-bPI-PLCs of the family (as outgroups) inferred a complex history (Fig. 8). The *Chalaropsis* and *B. basicola* Cer-PLCs were basal to those of *Ceratocystis*, consistent with the hypothesis that at least one Cer-PLC was present in the most recent common ancestor of the *Berk-Chal-Cer* Clade and that there were separate expansions of the Cer-PLCs in *Ceratocystis* and *Berkeleyomyces*.

**Figure 8.**
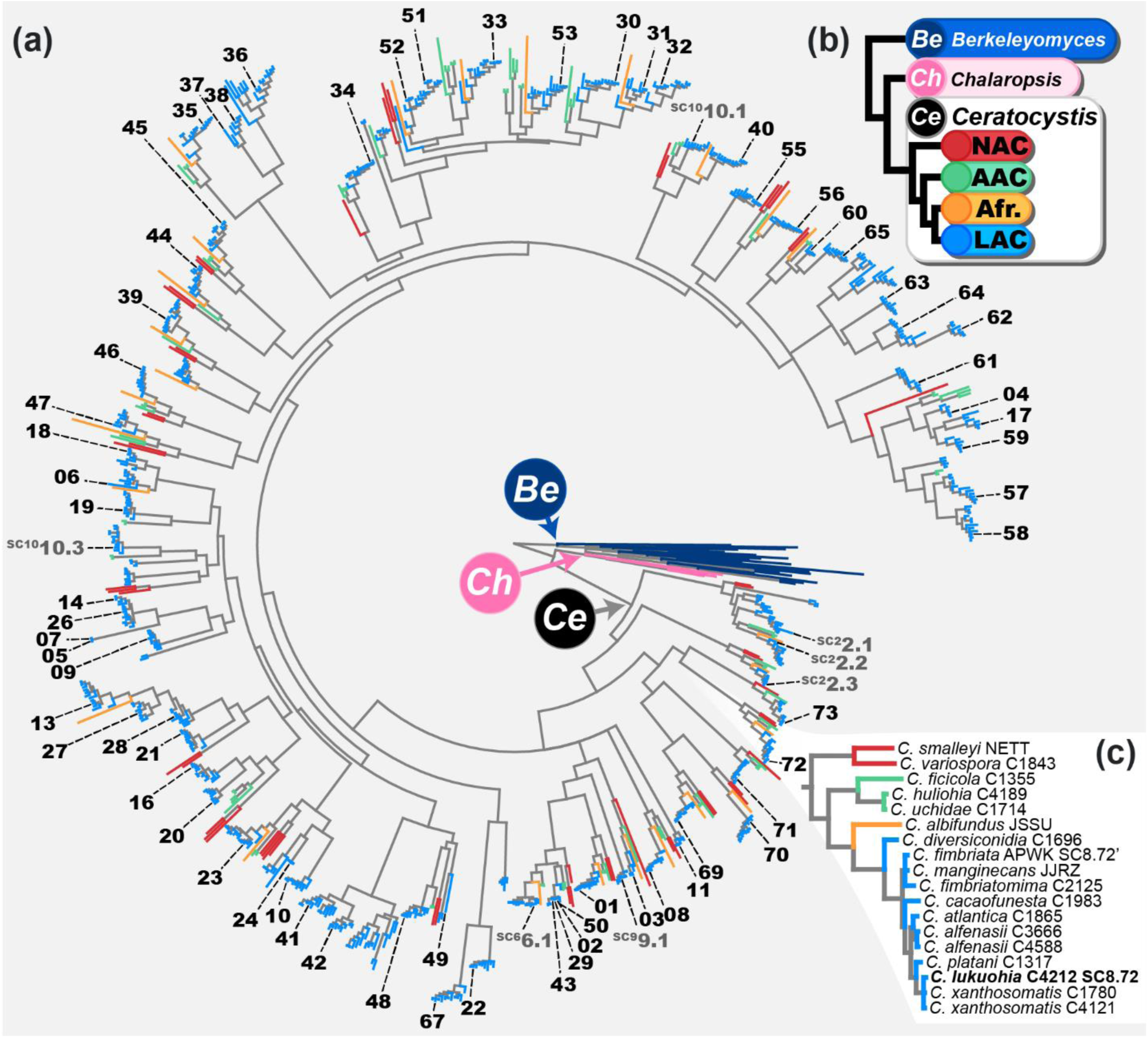
Circular tree produced from Bayesian analysis of more than 900 Cer-PLC genes (A). Colored branches indicate clade of source genome: *Berkeleyomyces* (dark blue), *Chalaropsis* (pink), and *Ceratocystis* species from the Latin American Clade (LAC; light blue), Asian-Australasian Clade (AAC; green), North American Clade (NAC; red) and African lineage (yellow). Labels indicate Cer-PLCs from *C. lukuohia*, with the numbers as defined in Fig 2 and Supplemental Table 4. (B) A simplified phylogram of the species relationship among these clades (modified from Fig. 5). (C) An expanded view of the clade containing *C. lukuohia* Cer-PLC 8.72, where all four geographic clades are present and with a topology as expected from the other gene trees.

Within *Ceratocystis*, Cer-PLCs grouped into many distinct clades, including at least 77 estimated orthologous gene clusters (OGCs) inferred from amino acid analyses. As suggested in the *C. lukuohia-C. fimbriata* analysis (Fig. 4), the clades containing the three Cer-PLCs on supercontig 2 of *C. lukuohia* (SC2.1, 2.2, and 2.3) were at the base of the Cer-PLC tree. The SC2 Cer-PLC subclades included all four geographic clades (Fig. 8), but *Ceratocystis* species outside of the LAC had only two of these basal SC2 Cer-PLCs.

The five Cer-PLCs at the 3’-end of the Cer-PLC cluster (SC8.69–SC8.73 in *C. lukuohia*) also were broadly conserved across the genus and formed an early diverging clade (Fig. 8). About half of the lineages of Cer-PLC cluster genes contained representatives of each of the geographic clades of *Ceratocystsis*, each lineage broadly following the topology of the species in the four-gene tree (Figs. 5 and 8B). Other lineages included only the Cer-PLCs of members of the LAC, apparently representing duplications and diversifications since the origin of the LAC. The sublineages that included the SC6, SC9 and SC10 Cer-PLCs of *C. lukuohia* each comprised Cer-PLCs of only LAC species (Fig. 8).

### Evolution of the Cer-PLC cluster

The hypothesis that the *Ceratocystis* Cer-PLC cluster developed by insertions of Cer-PLCs into an ancestral gene cluster was tested by annotating (at 50% nucleotide identity) 81 Microascales genomes with a database of the three 5′ flanking (a, b, c) and the three 3′ flanking (h, i, j) genes from the *C. lukuohia* cluster. Five of the six flanking genes were broadly present in the family and in the same orientation, but at low levels of homology. All genera of Ceratocystidaceae had at least one representative genome with orthologs of ‘b’, ‘h’, ‘i’, and ‘j’, and except for the Cer-PLCs of *Ceratocystis*, the spacing between the genes was similar to that of *C. lukuohia* (Fig. 3). The transcription activator ‘a’ was also found throughout the family, but with a sharply declining homology (by nucleotide identity) with greater distance from *Ceratocystis*. The hypothetical protein, ‘c’, appeared to be unique to *Berkeleyomyces, Chalaropsis*, and *Ceratocystis*. Some genomes of *Ambrosiella, Endoconidiophora,* and *Davidsoniella* had unique ORFs inserted into different places among the flanking genes. The Ext-bPI-PLCs found in *Thielaviopsis*, *Endoconidiophora*, *Davidsoniella*, and *Huntiella* (Fig. 6) were not found on the contigs with the flanking genes.

Only the *Ceratocystis* genomes had PLC-like genes in the purported ancestral cluster, which is best characterized by that of *Chalaropsis*, where the six flanking genes were contiguous within a compact cluster length of 17.7 kb (Fig. 3). The b-c gap was small (438 bp), as in the *C. lukuohia* cluster (456 bp). The c-h gap was also small (1,300 bp) with no intervening ORFs. Both gaps were present and similarly small in the other three *Chalaropsis* genomes, though in our C1375 assembly, one contig ended just after ‘h’ and another just before ‘j’. In all *Chalaropsis* genomes, the single Cer-PLC was on a separate contig.

The six flanking cluster genes were present in *Berkeleyomyces* on a single contig, in the same order and orientation as in *Chalaropsis*, but a large gap (12,169 bp) separated ‘b’ and ‘c’ and a smaller gap (1,848 bp) separated ‘c’ and ‘h’ (Fig. 3). Four ORFs in the larger b-c gap appeared to be unique to *Berkeleyomyces*. Products of two were identified via InterProScan as a metallo-beta-lactamase type 2-like protein (IPR050855) and a NMRA-like protein (IPR051609), and the others were a putative transposable element and a hypothetical protein. The cluster length (from beginning of transcription activator ‘a’ to the end of ‘j’) was 27.7 kb. The 25 Cer-PLCs were scattered across eight contigs, none of which contained the cluster flanking genes (Fig. 3).

A small number of *Ceratocystis* contigs ended immediately after the 5’ set of flanking genes or began immediately before the 3’ set of flanking genes, but almost all of the contigs with flanking genes included Cer-PLCs in an orientation consistent with the cluster arrangement seen in *C. lukuohia* (Fig. 3). The exceptional inversion previously noted in *C. fimbriata* and the unique HET gene (Fig. 2) were only seen in that species. The internal cluster genes in *C. lukuohia and C. fimbriata* (d, e, f) were unique to the LAC, but not all species in the clade had all three genes (Fig. 5B).

### Regulator genes and binding motifs in the Cer-PLC cluster

Fungal CeGAL transcription factors (also known as zinc cluster or Zn_2_Cys_6_ regulators), such as flanking gene ‘a’ (Fig. 2), often begin and coregulate gene clusters, and similar promoter regions are often found upstream of the regulator and regulated genes (Fernandes et al. 1998, Kennedy et al. 1999, MacPherson et al. 2006, Guo and Wang 2014, Mayer et al. 2023). We tested cluster-associated genes for promoter motifs that were enriched relative to the nuclear genomes of *Ch. thielavioides*, *C. lukuohia* and *B. basicola* using MEME.

One motif (*Ch. thielavioides* motif 2) was enriched upstream of all six flanking genes in *Ch. thielavioides*, and a similar motif (*C. lukuohia* motif 3) was enriched upstream of the six flanking genes and 71 of the internal (full or partial) Cer-PLC genes in the *C. lukuohia* cluster (Fig. 9).

**Figure 9.**
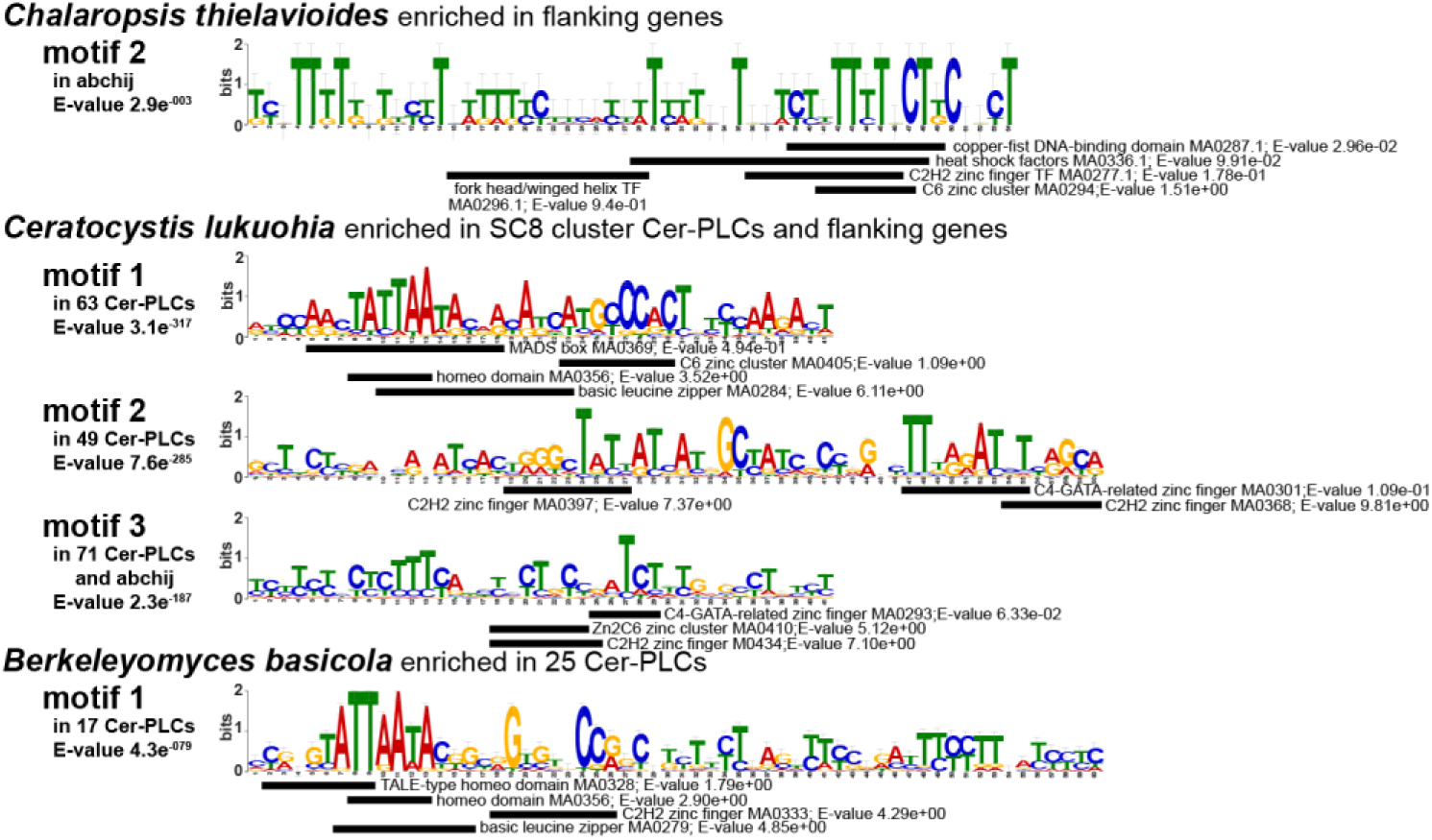
Motifs identified as enriched in the 1,000 base pairs (1 kb) preceding cluster genes and/or Cer-PLCs compared to other genes from the genomes of *Chalaropsis thielavioides, Ceratocystis lukuohia*, and *Berkeleyomyces basicola*, Sequence logos represent motifs as determined by MEME. Horizontal black bars below motifs indicate all hits (*Ceratocystis* and *Berkeleyomyces*) and the top five hits (*Chalaropsis*) for binding motifs in the JASPAR database as determined by TomTom. When these hits overlap, they are ordered from best (top) to worst (bottom) scoring hits by E-value.

Both motifs are CT-rich, typical of pyrimidine-rich regions of fungal promoter regions (Sibthorp et al. 2013, Sakekar et al. 2021), and both motifs contained typical CeGAL-type (Zn2C6) binding sites, suggesting that the CeGAL transcription factor ‘a’ was regulating the cluster in the most recent common ancestor of *Chalaropsis* and *Ceratocystis*.

Two additional high-scoring motifs (*C. lukuohia* motifs 1 and 2) were enriched before most of the *C. lukuohia* Cer-PLCs in the cluster but not before the Cer-PLCs found on other supercontigs (Fig. 9). The best-scoring of these (motif 1) included a CeGAL-type binding site typical of fungal gene clusters among possible hits, but it differed from *Ch. thielavioides* motif 2. It was found before transcription factor ‘a’ but not before the other flanking genes (Fig. 9), suggesting the development of a new CeGAL promoter region for the cluster after the integration of Cer-PLCs. *Ceratocystis lukuohia* motif 2 was only identified before the Cer-PLC genes and not before ‘a’ nor the other cluster genes. It apparently lacked a CeGAL-type binding site but had potential C2H2-type binding sites, suggesting another type of gene regulation.

No CeGAL-type cluster motif was identified among the cluster genes of *Berkeleyomyces* (Fig. 3). A single enriched motif was found before 17 of the 25 scattered *Berkeleyomyces* Cer-PLCs. Multiple transcription factor binding site types were predicted, and there were some similarities to the *Ceratocystis lukuohia* motif 1, but no CeGAL-type binding site was identified (Fig. 9).

## DISCUSSION

The proliferation of a bacterial-type phospholipase C gene (Cer-PLCs) in *Ceratocystis* represents one of the largest known expansions of a gene family in fungi. Large numbers of phosphatidylinositol-specific phospholipase C genes (PI-PLCs) were reported in *C. cacaofunesta* and some other *Ceratocystis* species in the LAC (Molano et al. 2018), and also in the North American species *C. destructans* (Maguvu et al. 2023). However, our examinations found that the PLCs lack critical residues that confer phosphatidylinositol specificity, and they represent a novel gene family that is unique to *Ceratocystis* and its closest relatives. The vast majority of *Ceratocystis* Cer-PLCs are found within what appears to be an ancestral gene cluster with a transcription regulator. The cluster was found to be dynamic and a general feature of the genus, with the aggressive plant pathogens in the LAC having the largest numbers of Cer-PLCs. Although the role of these novel Cer-PLCs in pathogenicity has not been demonstrated experimentally, their likely function in membrane disruption and their universal presence in a unique gene cluster strongly suggest their critical importance in the biology and evolution of *Ceratocystis*.

### Cer-PLCs as a novel enzyme family

Molano et al. (2018) reported that most of the *Ceratocystis* PI-PLCs had signal peptides, suggesting that they were secreted pathogenicity factors, and four of the enzymes were found in the secretome of *C. cacaofunesta* grown on artificial media. They concluded that *Ceratocystis* PI-PLCs maintain phosphatidylinositol specificity, though some of their proposed binding residues were inconsistent with the *B. cereu*s model for bPI-PLCs. We suggest that the active site of Cer-PLCs remains in the canonical position for bPI-PLCs, but active site residues have substantially changed, likely altering substrate specificity.

Bacterial-like PI-PLCs (bPI-PLCs; EC 4.6.1.13) are understudied in fungi but are distinct from eukaryotic PI-PLCs (ePI-PLCs; EC 3.1.4.11), which typically function in cell signaling. Prokaryotic bPI-PLCs are small, water-soluble, extracellular enzymes with a single functional domain (X domain) that cleaves phosphatidylinositol and related molecules in membranes. In bacteria, bPI-PLCs typically function as pathogenicity factors and are not known to be involved in signal transduction (Griffith and Ryan 1999). Fungal phospholipases have been reported to be potential pathogenicity determinants in diseases in plants and humans, including roles in signal transduction and adherence to or disruption of host membranes (Ghannoum 1998, 2000; Chayakulkeeree et al. 2008; Rho et al. 2009; Djordjevic 2010; Park et al. 2013; Barman et al. 2018; Xie et al. 2022; Li et al. 2025). However, such studies have either focused exclusively on ePI-PLCs, did not make a distinction between bPI-PLCs and ePI-PLCs, or involved other types of PLCs. *Magnaporthe oryzae* has multiple ePI-PLC genes (Rho et al. 2009; Barman et al. 2018). Secreted PI-PLCs have been reported in *Fusarium* and *Metarhizium* (Ma et al. 2010; Gao et al. 2011; Molano et al. 2018).

We found three types of bacterial-type PI-PLCs genes in the Ceratocystidaceae that were distinguished by the likely localization of their translated products. The cytoplasmic (Cyt-PI-PLCs) and vacuolar (Vac-PI-PLCs) bPI-PLC genes are each single-copy genes present throughout the Ceratocystidaceae and appear to have homologs in some related ascomycetes, suggesting they each arose prior to the origin of the family. These genes may not be relevant to *Ceratocystis* pathology as they are present in saprophytes, plant pathogens, and insect symbionts, and the Cyt-PI-PLC gene has been found in yeast. The translated products of the Cyt-PI-PLC and Vac-PI-PLC genes lack signal peptides, but their functional domains are similar to bPI-PLCs, with similar or identical amino acids at critical sites expected for bPI-PLCs found in bacteria.

The single copy Ext-PI-PLCs found in *Huntiella*, *Endoconidiophora*, *Davidsoniella* and *Thielaviopsis* encode proteins that have a N-terminal signal peptide, and their functional domains are most similar to those of Vac-PI-PLCs, which may also have a secretory pathway (Shoji et al. 2014). A change in function of a duplicate form of a Vac-PI-PLC to an Ext-bPI-PLC with a novel independent secretion signal would be consistent with a recognized route of neofunctionalization after gene duplication, which allows for the accumulation of mutations important to the new function without loss of the original function (Ohno 1970; Zhang 2003; Copley 2020; Birchler and Yang 2022).

In bacteria, five conserved catalytic amino acid residues are critical for bPI-PLCs to function as pathogenicity factors against host membranes or in escaping host vacuoles (Kuppe et al. 1989; Essen et al. 1996; Gässler et al. 1997; Moser et al. 1997; Hondal et al. 1998; Griffith and Ryan 1999; Kravchuk et al. 2001; Guo et al. 2008). Even single-nucleotide mutations at these positions can eliminate or drastically alter bPI-PLC activity (Gässler et al. 1997, Wei et al. 2005). In *Bacillus cereus*, these sites are numbered His32, Asp33, Arg69, His82, and Asp274, and each is highly conserved for the Cyt-PI-PLC, Vac-PI-PLCs and Ext-PI-PLCs of the Ceratocystidaceae. However, alternative substitutions were found in all of the studied Cer-PLC genes. The most dramatic change is at Arg69, where the positively charged Arg69 is replaced by the uncharged serine or threonine in nearly all Cer-PLCs. In bacteria, Arg69 positions the phosphate group for nucleophilic attack, and its replacement can alter substrate specificity (Kravchuk et al. 2001; Kravchuk et al. 2003; Wei et al. 2005). Additionally, nearly all Cer-PLCs replaced Lys115 (usually with glutamine), and the second histidine residue (His82) was conspicuously not conserved. In bacteria, a change of Lys115 to glutamine alters the pKa of His32 such that, when paired with an Arg69 substitution (to Asp), converts the enzyme from being PI-specific to a broad-target glucose phosphatase and reduces the absolute dependency on His82 (Heinz et al. 1995; Gässler et al. 1997; Feng et al. 2004). Mutation of His82 can also potentially allow for new substrates (Gässler et al. 1997). The retention of the essential catalytic His32 and histamine-positioning Asp33 and 274 (Heinz et al. 1995; Gässler et al. 1997) in most Cer-PLCs suggests that basic phospholipase C activity is maintained, even if substrate specificity is altered. We also noticed differences in the Cer-PLCs in the region of the “hydrophobic ridge,” which is a conserved region in the latter part of the bPI-PLC enzyme that is thought to mediate substrate binding, membrane specificity, and positioning of the catalytic residues after partial membrane penetration (Williams and Katan 1996; Katan and Williams 1997; Ellis et al. 1998; Griffith and Ryan 1999; Feng et al. 2003).

*Ceratocystis* PLCs were hypothesized to be both PI-specific and able to contribute to plant membrane disruption (Molano et al. 2018, Mora-Ocampo et al. 2021). However, phosphatidylinositol is only a minor component of plant cell membranes and contributes little to bulk architecture or integrity. Its importance has instead been associated with disruption of membrane-associated signaling (Cacas et al. 2016; Noack and Jaillais 2020). Mora-Ocampo et al. (2021) found elevated phospholipid metabolism in a susceptible cacao cultivar relative to a resistant cultivar inoculated with *C. cacaofunesta*, perhaps due to generalized phospholipid damage in the susceptible cacao genotype due to Cer-PLCs.

The unique Cer-PLCs of the *Berk-Char-Cer* Clade may have evolved from the Ext-PI-PLCs found in its sister clade (the *Thiel-Endo-David* Clade) following the Innovation-Amplification-Divergence (IAD) model, as exemplified in the MALS gene family expansion in yeast (Voordeckers et al. 2012). In IAD, a pre-existing promiscuous enzyme or a protein with weak, secondary activity becomes selectively advantageous, it is amplified through gene duplication and then diverges as paralogs specialize (Bergthorsson et al. 2007; Näsvall et al. 2012). The replacement of Arg69, a position critical for phosphate positioning (Kravchuk et al. 2001, 2003), by Ser/Thr in nearly all Cer-PLCs parallels this type of active-site changes that redirect catalysis toward novel substrates (Copley 2020).

The hypothesized evolution of Ext-PI-PLCs to Cer-PLCs largely tracks the phylogeny of the Ceratocystidaceae and their increasing pathogenicity. The single-copy Ext-PI-PLCs are found in genera that include weakly pathogenic wound colonizers (*Huntiella*), wood-staining fungi (*Endoconidiophora*), and minor or host-specialized plant pathogens (*Davidsoniella* and *Thielaviopsis*) that cause minor root diseases and fruit rots, with a few causing lethal diseases of trees (Harrington 2009, deBeer et al. 2014). The unusual aquatic *Seychellomyces sinensis* occupies an informative phylogenetic position between the Ext-PI-PLC genera and the Cer-PLC genera (Qiao et al. 2019), but a genome was not available for study. *Chalaropsis* species, which have a single Cer-PLC, are relatively minor root pathogens (Nag Raj and Kendrick 1975, Szabo and Harrington 2004, de Beer et al. 2014). *Berkeleyomyces basicola* has 25 Cer-PLC genes; it causes black root rot on a wide range of hosts, including substantial losses to cotton and other crops, typically involving black staining of host tissues (Nel et al. 2019).

The occurrence of large numbers of Cer-PLCs in a tight gene cluster appears to be the defining characteristic of the genus *Ceratocystis*. All *Ceratocystis* species are plant pathogens, often with very broad host ranges. They vary greatly in their aggressiveness to their various hosts, ranging from black rotters of storage roots and corms to colonizers of fresh wounds on woody hosts to systemic vascular wilt pathogens that are capable of killing large trees (Johnson et al. 2005, Thorpe et al. 2005; Harrington 2013, 2024; deBeer et al. 2014, Hughes et al. 2020). In most cases, a conspicuous brown to black staining of xylem tissues is evident, sometimes accompanied by necrosis of the secondary phloem and canker formation.

### The origin of the Cer-PLC gene cluster

While *Chalaropsis* species have only a single Cer-PLC gene, there were two separate and dramatic amplifications of Cer-PLCs in *Berkeleyomyces* and *Ceratocystis*. In *Ceratocystis,* most of the Cer-PLCs were found in a large gene cluster with a CeGAL-type transcription factor, which was also present in *Chalaropsis* and *Berkeleyomyces* but without Cer-PLC genes. In *B. basicola*, at least 25 Cer-PLCs are scattered across the genome, but they were not on the same contig as the CeGAL-type factor, and most had an apparent non-CeGAL transcription factor binding site.

The Cer-PLC gene cluster in *Ceratocystis* apparently arose through modification of an ancestral gene cluster comprising six genes in an arrangement (5’-abchij-3’) found in some form throughout much of the Ceratocystidaceae. The likely ancestral cluster is represented by the *Chalaropsis* cluster, which begins with genes coding for two nucleus-located proteins: ‘a’, a CeGAL Zn2-C6 transcription factor (Mayer et al. 2023) and ‘b’, an AAA+ ATPase, which may serve as a secondary regulatory transcription factor (Heallen et al. 2008; Fu et al. 2023). The downstream genes code for a hypothetical protein ‘c’ (which was only found in the *Berk-Chal*-*Cer* Clade), a membrane trafficking protein ‘h’ (a Rab-like, small GTPase), an energy metabolism protein ‘i’ (ATP synthase regulation protein nca2, Camougrand et al. 1995), and a lipid metabolism enzyme ‘j’ (a fatty acid hydrolase, likely present in the endoplasmic reticulum).

Upstream of each of the six flanking genes in *Chalaropsis* and *Ceratocystis lukuohia*, there were similar CT-rich motifs typical for binding sites of fungal promoters (Sibthorp et al. 2013; Sakekar et al. 2021) and consistent for a CeGAL (Zn_2_-C_6_) transcription factor. In *C. lukuohia*, the putative CeGAL binding site was also seen in front of most of the Cer-PLC genes. There was a second motif specific to the upstream promoter region of most Cer-PLCs and the CeGAL-type transcription factor gene ‘a’, but the CeGAL-type binding motif was not identified in the other flanking genes. This and other regulatory elements may have evolved alongside the expanding gene family, reminiscent of how effector genes in *Ceratocystis* and other fungi acquire or modify cis-regulatory elements during or after gene expansion (Richards et al. 2019; Fourie et al. 2020).

Gene clusters in fungi typically contain up to a few dozen genes encoding different enzymes in a shared biosynthetic pathway (Yu et al. 2004; Zeng et al. 2018). Nearly half of the characterized biosynthetic gene clusters in fungi include an adjacent pathway-specific transcription factor, often of the CeGAL type (Keller 2019; Meyer et al. 2023). The ancestral gene cluster found in *Chalaropsis* was likely coordinating genes related to energy metabolism, membrane trafficking, and lipid metabolism. The *Ceratocystis* Cer-PLC cluster of secreted and potentially membrane-disrupting enzymes may have developed through insertion of one or more Cer-PLCs between ‘c’ and ‘h’ of a gene cluster that was already under regulation by a CeGAL transcription factor.

Coordinated deployment of pathogenicity factors is a hallmark of aggressive fungal plant pathogens (Richards et al. 2019; Seong and Krasileva 2023). Gene clusters regulated by Zn₂Cys₆/CeGAL transcription factors may act as direct coordinators of enzyme secretion during pathogen attack (Rybak et al. 2017; John et al. 2024; Zhang et al. 2025), and in *Ceratocystis*, this would avoid constitutive production of the large arsenal of Cer-PLCs. Although Syazwan et al. (2025) did not list PLCs among the upregulated proteins produced by *C. manginecans* within the first 24 h of stem infection, four Cer-PLCs were detected in the secretome of *C. cacaofunesta* grown in a xylem medium at 72 h (Molano et al. 2018). Delayed expression after initial infection may be consistent with an inducible expression during active tissue colonization. However, the regulation of Cer-PLC production and secretion in *Ceratocystis* needs further study.

Among all the *Ceratocystis* genomes examined, only the sweet potato strain of *C. fimbriata* showed deviation in the arrangement of the cluster flanking genes. An inversion near the 5’ end of the cluster and the addition of a unique heterokaryon compatibility (HET) gene were found in all the genome assemblies of this species. Such HET-E proteins govern heterokaryon compatibility and may inhibit sexual compatibility with other strains (Daskalov et al. 2023, Espagne et al. 2002). Sweet potato strains of *C. fimbriata* are less sexually compatible with other strains of *Ceratocystis* in mating studies (Harrington et al. 2024), and the unique HET gene in this host-specialized strain (Baker et al. 2003; Li et al. 2016) may limit inter-strain interactions and lead to stability of its Cer-PLC cluster.

### Expansion of the Cer-PLC family in *Ceratocystis*

Phylogenetic analyses of *Ceratocystis* Cer-PLCs showed early divergence of the supercontig 2 (SC2) Cer-PLCs of *C. lukuohia* and *C. fimbriata* from the Cer-PLCs in the gene cluster on SC8. Across the genus, it appears that there are two major lineages of SC2-type Cer-PLCs, each of which have C-terminal extensions beyond the X domain: a metallo-hydrolase/oxidoreductase domain or a proline-rich intrinsically disordered region that may facilitate membrane integration and enhance PLC activity (Theillet et al. 2013; van der Lee et al. 2014). Some of the genomes of LAC species showed two tightly-linked and related SC2 Cer-PLCs with proline-rich extensions, indicating recent duplication. Both the phylogenetic analyses of the X domains and the unique C-terminal extensions suggest that the SC2 Cer-PLCs evolved separately from the Cer-PLCs in the gene cluster and may have been present before the Cer-PLC gene cluster developed.

Insertion and deletion of short syntenic segments of one to six Cer-PLC genes and their intergenic regions were evident in comparisons of Cer-PLC clusters. The most conserved segment is found at the 3’ end of the cluster, before ‘h’, where five closely related Cer-PLCs were found in the LAC species *C. lukuohia* (SC8.69 to 8.73) and *C. fimbriata*, as well as in the much smaller cluster (282,248 bp) of the recently released long-read genome assembly of *C. huliohia,* an AAC species (Nakamoto et al. 2026). These five tightly-linked and highly-conserved Cer-PLCs formed an early diverging clade of Cer-PLCs, suggesting that the initial insertion and expansion of Cer-PLCs began with one or more of these Cer-PLCs. Later duplications appeared more haphazard. Some conserved Cer-PLC genes that were found in all geographic clades of *Ceratocystis* were scattered across the gene cluster, with their position within the cluster showing little resemblance to their position in the Cer-PLC tree. In several cases, pseudogenes were identified that lacked some of the 5′ signal peptide portion of the gene or some of the 3′ functional domain. Also noteworthy, non-cluster Cer-PLCs on other supercontigs (on SC6, SC9, and SC10 in *C. lukuohia*) were phylogenetically nested within Cer-PLCs found within the gene cluster, suggesting that they had escaped the cluster.

The precise mechanisms of gene duplication (Welch et al. 1990; Jelesko et al. 1999; Zhao et al. 2014) are unclear, but the patterns of truncated genes and large indels in a tight gene cluster suggest frequent recombination events due to repetitive elements. In crosses between strains, misalignment of elements of a multi-copy gene family may lead to unequal and double crossover events (Mehradi et al. 2017), which could lead to insertion or deletion of large syntenic segments, highly variable copy numbers, chimeric Cer-PLCs, and pseudogenes. Reiterative unequal crossover events are thought to contribute to rapid amplification of tandemly arrayed rDNA genes in fungi (Smith 1976; Szostak & Wu 1980; Salim & Gerton 2019; Lofgren et al. 2019), and frequent recombination among copies of the rDNA array has been noted in *Ceratocystis* (Harrington et al. 2014, 2024). The Cer-PLCs are not tandem repeats, but even small regions of similar Cer-PLC genes would provide for sufficient non-homologous pairing and recombination hot spots (Jinks-Robertson et al. 1993).

As an alternative type of repetitive element, transposable elements are thought to commonly contribute to genome rearrangements and expansion in fungi through transposon-mediated duplication (Stajich 2017, Mehradi et al 2017). Some plant-pathogenic fungi exhibit copy-number variation apparently facilitated by a “two-speed genome” architecture (Raffaele and Kamoun 2012; Dong et al. 2015), such as lineage-specific virulence-related genes in *Verticillium* that are clustered within rapidly evolving, transposon-rich regions of the genome (Faino et al. 2016). However, the Cer-PLC cluster is not particularly transposon-rich (Torres et al. 2020).

The degree of expansion of the Cer-PLC gene family within a single genetic region appears unprecedented in fungi, with the exception of ribosomal DNA operons (Lofgren et al. 2019). In the microsporidian *Nematocida displodere*, there was a significant pathogen-host driven expansion of a genus-specific gene family, but the genes were not in a cluster (Reinke et al. 2017). The CBM18 domain expansion in *Batrachochytrium dendrobatidis* has 65–90 domain copies, but the domains are distributed across multiple proteins (Abramayan and Stajich 2012; Farrer et al. 2017; Stajich 2017). The M36 metalloprotease virulence genes in *B. salamandrivorans* have 177 copies, but they are distributed across multiple repeat-rich compartments (Wacker et al. 2023).

Outside of fungi, some of the most extreme expansions of genes occur in systems where pathogen-derived factors interface directly with host cellular components (Lažetić and Troemel 2021). In response to pathogen pressure, nucleotide-binding and leucine-rich disease resistance genes in plants may form large, rapidly evolving tandem clusters that vary dramatically in copy number among lineages (Meyers et al. 1998; van Wersch and Li 2019), diversifying through unequal crossover and gene conversion (Meyers et al. 1998, 2003; Baggs et al. 2017).

Rattlesnake venom metalloproteinase (SVMP) genes expanded from a single ancestral locus into a tandem array of around 30 genes within a 1.3 Mb region, driven by sequential duplication, intragenic deletion, and neofunctionalization (Giorgianni et al. 2020). In vertebrates, olfactory receptor genes (OR) form the largest known multigene superfamily, containing hundreds of genes scattered across the genome, generated through repeated tandem duplication and diversification (Glusman et al. 2001; Niimura and Nei 2003). These genes follow a birth-and-death model of evolution in which new genes arise by duplication, diverge in sequence and function (neofunctionalization), and are pseudogenized or retained depending on selection pressure (Nei and Rooney 2005; Nei et al. 2008). Although the Cer-PLC expansion in *Ceratocystis* has largely taken place in a regulated gene cluster, the fundamental drivers of gene expansion by unequal crossing over and lineage-specific diversification under selection may be similar to the examples above.

### Expansions of Genomes, the Cer-PLC cluster and the LAC

Genomes of the Ceratocystidaceae are relatively small, fitting the general trend for insect-associated fungi, although the increases in genome size with pathogenicity in the *Berk-Chal-Cer* Clade are not necessarily general trends across Sordariomycetes (Fijarczyk et al. 2025).

*Chalaropsis* had the smallest estimated genome sizes of the family (23.2–23.9 Mb) and the smallest number of Augustus-estimated genes (6626). In the study of 552 genomes of Sordariomycetes by Fijarczyk et al. (2025), only *Ceratocystiopsis*, *Ophiocordyceps*, and *Tolypocladium* had smaller genomes than *Chalaropsis,* and only *Ambrosiella* had fewer predicted genes. A contraction in genome size in a *Chalaropsis*-like ancestor would suggest expansions in *Berkeleyomyces* (24.2–25.5 Mb, 7145 genes) and *Ceratocystis* (25.8–33.2 Mb, 7210 genes) that are much larger than the simple expansion of 24 Cer-PLCs in *Berkeleyomyces* and the 282 Kb and 530–543 Kb lengths of the Cer-PLC clusters in the AAC (*C. huliohia*) and the LAC (*C. fimbriata* and *C. lukuohia*). The largest and most variable genome sizes were seen in the LAC, with an average of 7382 estimated genes vs. 6953 in the other *Ceratocystis* species, a difference much greater than the differences in Cer-PLC copy number alone (51–92 vs 26–36 in the other *Ceratocystis* species).

The dynamic nature of chromosome rearrangements in the LAC (Fernandes et al. 2022) and its genome expansion, especially in the Cer-PLC cluster, suggest a major evolutionary event, perhaps a hybridization, at the origin of the clade. *Ceratocystis* species are capable of selfing through unidirectional mating type switching or crossing with what could be considered other species (Ferreira et al. 2010; Harrington et al. 2014, 2024; Barnes et al. 2016; Li et al. 2016; Oliveira et al. 2018; Fourie et al. 2018; Kanzi et al. 2020; Van der Walt et al. 2023).

Mitochondrial introns can be vegetatively transmitted between even distantly-related species of *Ceratocystis* (Mayers et al 2023). The global movement of infected plant propagation material allows allopatric strains or species to hybridize and potentially produce novel combinations of Cer-PLCs, which may degrade different membrane components of different hosts, perhaps contributing to wide variation in host-range and aggressiveness (Baker et al 2003; Harrington et al. 2011; 2014; 2024).

Expanded gene families may be most pronounced in the most ecologically dominant or pathogenically aggressive (ecologically successful) lineages (Lažetić and Troemel 2021). The genus *Ceratocystis* is thought to be relatively young (∼15 MYA, Mayers et al. 2020), and the four-gene tree suggested that the LAC is the youngest clade. The LAC shows almost no morphological variation (Lynn et al. 2026), yet it is the most species-rich clade of *Ceratocystis* and the most ecologically and economically destructive clade (Harrington et al 2024). The species in the LAC generally have wide host ranges, but some are apparently host-specialized. The host-specialized *C. platani*, the *Syngonium* strain of *C. xanthosomatica* and the sweet potato strain of *C. fimbriata* (Baker et al. 2003, Thorpe et al. 2005) have somewhat lower numbers of Cer-PLCs compared to other LAC species, perhaps because there is less need for a diversity of Cer-PLCs for a narrow host range.

The greatest number of Cer-PLCs (64–92) and the widest host ranges are found in the South American species, especially *C. manginecans,* strains of which have very broad and variable host ranges in pathogenicity tests (Baker et al. 2003; Harrington et al 2011; Harrington et al. 2024).

Higher copy numbers of Cer-PLCs were mostly found in aggressive strains in geographic regions where the pathogen had been introduced (Harrington et al. 2024). While the introduced strain of *C. manginecans* in Oman had the highest number of Cer-PLCs (92), only 64 Cer-PLCs were identified in the host-specialized, seedling-blight strain on *Carapa*, a host tree native to the Amazon Basin that is attacked by an apparently native population of *C. manginecans* (Valdetaro et al. 2019). The spreading epidemic of *C. manginecans* from Oman to China to Southeast Asia to Palau on a wide range of hosts appears to be due to two or more introduced strains from South America that have hybridized (Harrington et al. 2024, Johnson et al. 2025, Lynn et al. 2026). Recombination within the Cer-PLC cluster has not been studied, and changes in phenotype have not yet been associated with variation in Cer-PLCs. However, the Cer-PLC cluster may well be responsible for the highly dynamic host range of *C. manginecans* and may lead to the emergence of new super pathogens as strains continue to be dispersed in infected, vegetatively-propagated plant material (Harrington et al. 2024).

## Acknowledgements

The laboratory assistance of Jenna Vickery and Grace Schulte is gratefully acknowledged.

## Data Availability Statement

All studied genome assemblies are publicly available, and the accession numbers are listed in Table 1. Supplementary tables 4 and 5 describe the locations of identified phospholipase and related genes in the genomes of *Ceratocystis lukuohia* and *C. fimbriata*.

---

Supplemental Table 1. *Ceratocystis lukuohia* genome assemblies, contig characteristics and BUSCO analysis using different read lengths and assemblers.

Supplemental Table 2. Size, presence of telomeres, and number of predicted transposable elements and genes on 12 supercontigs representing the nuclear genome of *Ceratocystis lukuohia*.

Supplemental Table 3. Sizes, contigs and predicted genes in the draft genome assemblies of three *Ceratocystis* species.

Supplemental Table 4. Distribution and target localization of 88 predicted phosphatidylinositol-specific phospholipase-C (PI-PLC) genes on the 12 supercontigs of the *Ceratocystis lukuohia* genome.

Supplemental Table 5. Distribution and target localization of 77 predicted phosphatidylinositol-specific phospholipase-C (PI-PLC) genes on the 9 supercontigs of the *Ceratocystis fimbriata* genome.

Supplemental Table 6. Non-PLC genes flanking and internal to the Cer-PLC cluster in *Chalaropsis*, *Ceratocystis*, and *Berkeleyomyces*.

**Supplemental Figure 1.**
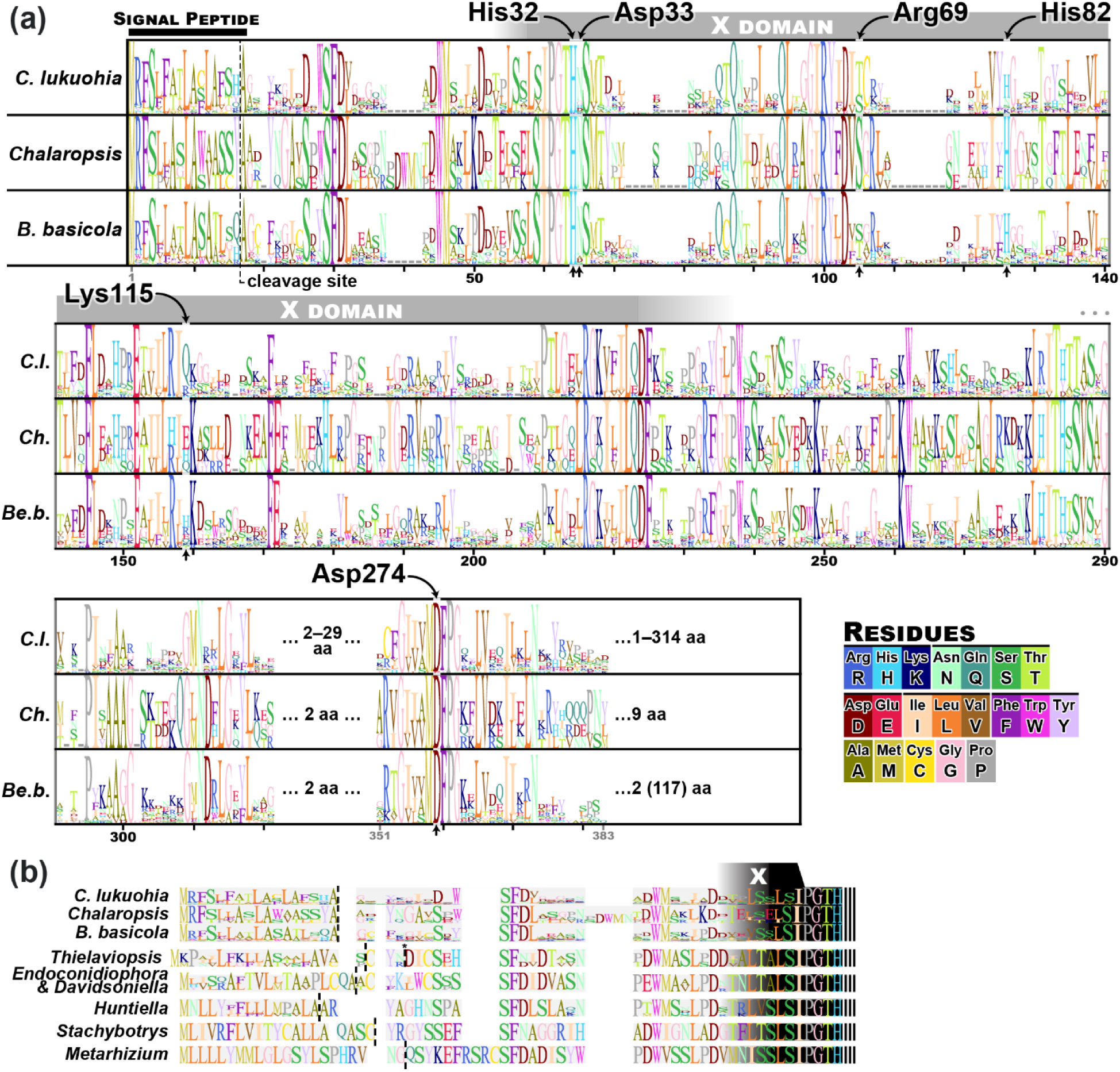
(A) Sequence logos of the amino acid conservation among Cer-PLCs in an alignment of Cer-PLCs from *C. lukuohia* (75 Cer-PLCs from isolate C4212), *Chalaropsis* (one Cer-PLC each from the four genomes used in this study), and *Berkeleyomyces basicola* (25 Cer-PLCs from PJAC00000000). Scale at bottom indicates amino acid position in the combined alignment. A black horizontal bar indicates the signal peptide, a dotted vertical line the cleavage site, and a grey box the approximate location of the X domain. Important residues discussed in the text are labelled. (B) Sequence logos of predicted signal peptide regions (and beginning of X domain) for selected Cer-PLCs, with predicted cleavage sites (per SignalP) indicated by dotted vertical lines. Residue logos are proportional to the percentage occurrence of that residue at that site, colored as per the legend.

**Supplemental Figure 2.** Original raw tree used to construct Figure 6. Branch support values are Bayesian posterior probability values rounded to four significant digits. Scale bar = 0.4 estimated substitutions per site.

