## Supplementary figures and images for "Evolution of a large and diverse phospholipase gene cluster that defines the plant pathogenic genus *Ceratocystis*"

### Supplemental Figure 1

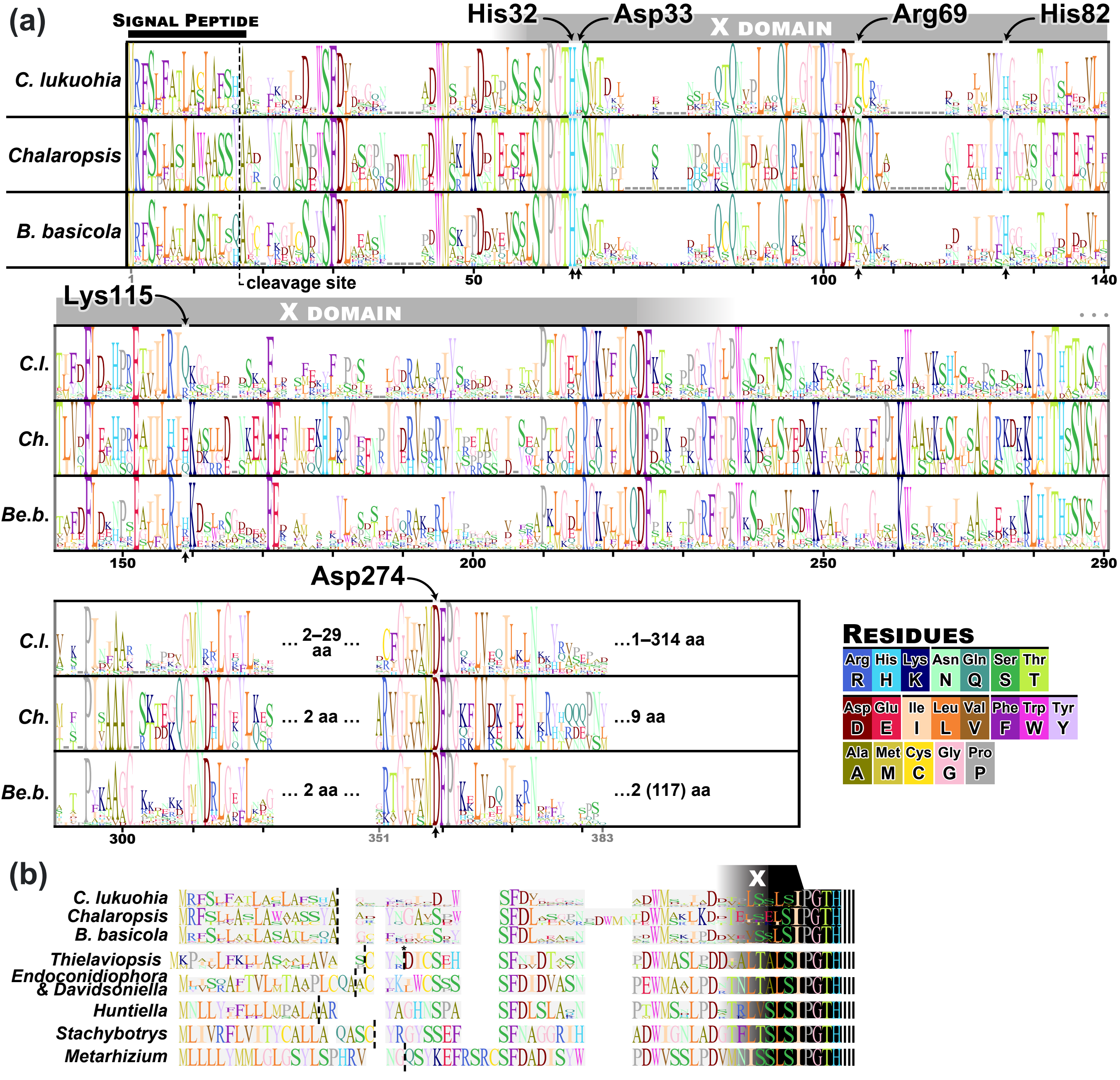

### Supplemental Figure 2

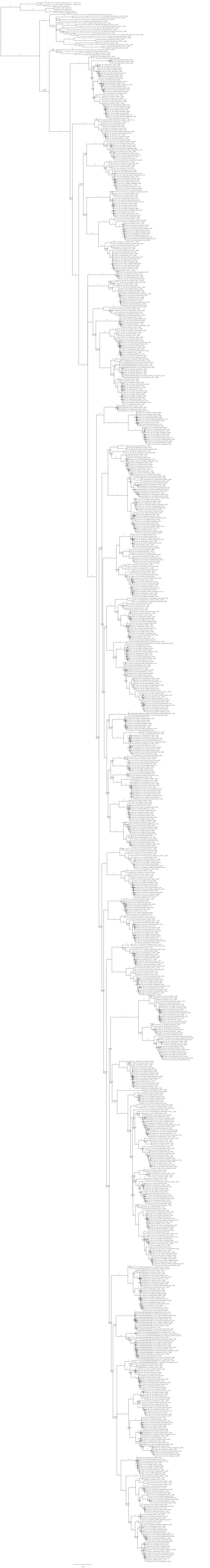
